# The Human Cerebello-Hippocampal Circuit Across Adulthood

**DOI:** 10.1101/2025.02.17.638640

**Authors:** Tracey H. Hicks, Thamires N. C. Magalhães, Jessica A. Bernard

**Author notes:** Corresponding Author: Tracey Hicks, M.A. Department of Psychological and Brain Sciences Texas A&M University 4235 TAMU, College Station, TX 77840 (979)845-2581. Co-author emails: Thamires N. C. Magalhães Jessica A. Bernard.

## Abstract

Direct communication between the hippocampus and cerebellum has been shown via coactivation and synchronized neuronal oscillations in animal models. Further, this novel cerebello-hippocampal circuit may be impacted by sex steroid hormones. The cerebellum and hippocampus are dense with estradiol and progesterone receptors relative to other brain regions. Females experience up to a 90% decrease in ovarian estradiol production after the menopausal transition. Postmenopausal women show lower cerebello-cortical and intracerebellar FC compared to reproductive aged females. Sex hormones are established modulators of both memory function and synaptic organization in the hippocampus in non-human animal studies. However, investigation of the cerebello-hippocampal (CB-HP) circuit has been limited to animal studies and small homogeneous samples of young adults as it relates to spatial navigation. Here, we investigate the CB-HP circuit in 138 adult humans (53% female) from 35-86 years of age, to define its FC patterns, and investigate its associations with behavior, hormone levels, and sex differences therein. We established robust FC patterns between the CB and HP in this sample. We predicted and found negative relationships between age and CB-HP FC. As expected, estradiol levels exhibited positive relationships with CB-HP. We found lower CB-HP FC with higher levels of progesterone. We provide the first characterization of the CB-HP circuit across middle and older adulthood and demonstrate that connectivity is sensitive to sex steroid hormone levels. This work provides the first clear CB-HP circuit mapping in the human brain and serves as a foundation for future work in neurological and psychiatric diseases.

## Introduction

Both the hippocampus and cerebellum exhibit atrophy in aging, play a role in cognition, and are modulated by sex steroid hormones^1–16^. Importantly, these regions have shown coactivation, synchronized neuronal oscillations, and bidirectional relationships on a cellular level^17^, suggesting direct communication between these regions. Further support for structural and functional connectivity between these regions has been shown in animal studies. Rabies virus tract tracing in rodents identified three primary pathways linking the cerebellum to the hippocampus^18^, while optogenetic stimulation of the cerebellum impacts hippocampal function and hippocampal-mediated behavior^19^. In humans, the hippocampus and cerebellum coactivate during tasks requiring spatio-temporal prediction of movements in visuomotor integration^20^. While this has clear implications for both spatial and cognitive-motor abilities, it may also be applied to episodic memory^17^. That is, much like spatial representations of places, routes, and maps, our memories are created in time-centered sequences resembling a conceptual path or route marked by life events^21^. As both cognitive and motor abilities decline with age^22–24^, interactions between these regions may be especially important. However, investigation of the cerebello-hippocampal (CB-HP) circuit has been limited to animal studies^18,19,25^ and small homogeneous samples of young adults as it relates to visuospatial abilities^20,26,27^.

Evidence suggests that the cerebello-hippocampal circuit may be impacted by sex steroid hormones. Higher levels of estradiol have been linked to greater functional network coherence and greater cortical to subcortical functional connectivity (FC) at rest^28–31^, whereas higher progesterone has displayed mixed relationships of higher and lower cortical to subcortical and intracortical FC^29,30,32–35^. Importantly, both the cerebellum and hippocampus are dense with estradiol and progesterone receptors relative to other brain regions^36^. Sex hormones are established modulators of both memory function and synaptic organization in the hippocampus in non-human animal studies^9^. In women, lower levels of sex hormones during the menopausal transition have been associated with altered hippocampal activity during memory tasks^7,8^. Further, postmenopausal women show lower cerebello-cortical and intracerebellar FC compared to reproductive aged females^2^.

Broadly, aging has been linked to deficits or declines in cognitive performance^22,37^, functional brain changes^38–42^, and a drop in sex hormones^29,43,44^. Functionally, aging is associated with reduced network segregation and increased integration of brain networks that have been linked to poorer cognitive performance^38–40,42,45^. Regarding sex hormones, females experience up to a 90% decrease in ovarian estradiol production after the menopausal transition^46^. The significant fluctuations and inevitable drop in sex steroid hormones during the menopausal transition and post - menopause have been associated with cognitive decline in females^47–50^. Further, higher levels of endogenous estradiol have been associated with better working memory performance^51,52^. Given the broad age-related impacts on brain FC, hormones, and cognition, coupled with the role of both the CB and HP in cognitive performance, exploration of this circuit in the context of these factors stands to be particularly informative for our understanding of aging and preclinical indicators of pathological decline.

Here, we first seek to quantify CB-HP interactions across middle and older adulthood. Second, we will investigate the relationship between the CB-HP circuit and both behavioral performance and sex steroid hormone levels. We will also investigate sex differences in this circuit. More specifically, we hypothesized that age would negatively correlate with CB-HP connectivity. Further, we predicted that better behavioral performance would be associated with higher CB-HP FC. We further expected higher CB-HP connectivity with greater 17β-estradiol levels^28–31^. As previous findings with higher levels of progesterone and FC have been mixed ^29,30,32–35^, we did not have specific predictions for CB-HP connectivity with progesterone levels.

## Methods

### 1.1. Study sample

Participants (total *n*=175) were enrolled as part of a larger study on aging. All participants underwent a battery of cognitive and motor tasks and saliva samples were collected for sex-steroid hormone quantification during this assessment (described below). After the behavioral visit the participants returned for a magnetic resonance imaging (MRI) session approximately two weeks later. However, due to unexpected delays related to the Covid-19 pandemic, the time between the two sessions ranged from 0 to 216 days between participants with a mean of 34.08 days and standard deviation of 40.22 days in our sample. Here, we focused on salivary hormone assays, behavioral performance, and relationships with brain imaging data.

Exclusion criteria were history of neurological disease, stroke, or formal diagnosis of psychiatric illness (e.g., depression or anxiety), contraindications for the brain imaging environment, diagnosis of mild cognitive impairment or dementia, and use of hormone therapy (HTh) or hormonal contraceptives (intrauterine device (IUD), possible use of continuous birth control (oral), and no history of hysterectomy in the past 10 years. For our analyses here we focused only on those with available neuroimaging, hormone level, and cognitive performance data. Thus, our final sample included 138 participants (74 females (*M* = 56.73 years, SD = 12.53) and 64 males (*M* = 56.58 years, SD = 12.53); range across both groups, 35-86 years); however, our sample size varies slightly across analyses based on participants with usable hormone data. Regarding ethnicity, our sample was 88.4% Caucasian and 11.6% Hispanic/Latino (**Supplementary Table 1**). Demographic details are presented for generalizability purposes and we recognize that these are social and political categories given meaning by social, historical, and political forces^53^. Race and ethnicity will not be evaluated separately in this study, as information about socio-economic status is better indicated for causal inferences in neuroimaging^53^ and that information was not collected in this study.

All study procedures were approved by the Institutional Review Board at Texas A&M University, and written informed consent was obtained from each participant prior to initiating any data collection.

### 1.2. Behavioral Testing

The behavioral testing sections (section 1.2. through 1.5.) utilize standardized text to ensure consistency and reproducibility in reporting the research protocol across work associated with this study^54–57^. Participants completed a battery of cognitive and motor tasks to quantify attention, processing speed, working memory, delayed recall, executive functioning, grip strength, fine motor abilities, and cognitive-motor integration. A commonly used screening tool that assesses global cognitive functioning, the Montreal Cognitive Assessment (MoCA) was also included^58^. The Wechsler Adult Intelligence Scale, 4th Edition (WAIS-IV) Letter-Number Sequencing subtest^59^ broadly assessed working memory. The Wechsler Memory Scale, 4^th^ Edition (WMS-IV) Symbol Span subtest was used to assess visual-spatial working memory^60^. Episodic memory was assessed via the Shopping List Memory Task^61^ which was administered on a computer.

#### 1.2.1. The Stroop Task

The Stroop Task^62^ was adapted to computer format and was used to gauge executive function. Both congruent and incongruent items were displayed for 2.6 seconds each. There were 10 blocks with untimed (self-paced) breaks in between each block. This study specifically examined the Stroop effect as an outcome. Mathematically, the score was the difference in reaction times between congruent (e.g., the word “blue” in blue font) and incongruent (e.g., the word “blue” in red font) items (incongruent – congruent). A higher score indicated a slower reaction time on incongruent trials, score of 0 indicated performing the same (on average) for both congruent and incongruent trials, and a negative score indicated a faster reaction time on incongruent trials. Those with incomplete data (n = 1; 9/10 blocks incomplete) and outliers with values greater than 3 standard deviations from the mean (n=1) were also excluded from calculations for this task.

#### 1.2.2. Sequence Learning Paradigm

An explicit sequence learning paradigm was based on a task created by Bo and colleagues^63^. This task assessed learning of motor behaviors and working memory. The task consisted of 6 random blocks with 18 items each interspersed with 9 sequence blocks with 36 items each. In sequence blocks, participants learned a 12-element sequence shown 1 second per element and repeated this sequence 4 times in each block. In the random blocks, participants learned a different sequence each time. For both blocks, they were shown a sequence of filled squares and asked to repeat it in its entirety. We chose to quantify learning by calculating the slope of task accuracy (labeled as sequence learning accuracy in the results section) from Block 1 to Block 9. Mathematically, the slope of task accuracy value was derived using the slope of a regression line equation to evaluate change across trials: m=(n(∑xy) − (∑x)(∑y))/ (n(∑x^2^) − (∑x)^2^). Where ‘∑x’ was the sum of trial numbers, ‘∑x^2^’ was the sum of squares of trial numbers, ‘∑y’ is the sum of performance accuracies, and ‘∑xy’ is the sum of the corresponding trial numbers and their accuracies. Conceptually, we would expect a positive slope for accuracy if learning/improvement had occurred across trials and a negative or no slope if it had not (i.e., a score above 0 indicated learning, with higher scores indicating a higher learning curve). To statistically evaluate whether learning had occurred, we conducted a one-sample t-test to determine if the Sequence learning slope was significantly different from zero.

#### 1.2.3. Purdue Pegboard Assembly

Cognitive-motor abilities were assessed via the Purdue Pegboard Task^64^. The Purdue Pegboard is comprised of four subtests: dominant hand, non-dominant hand, both hands simultaneously, and assembly task. We focused our investigations on the assembly subtest due to the integration of cognitive and motor demands of the task which is particularly relevant to our investigation of the cerebellum and cerebello-hippocampal connectivity. This subtest required participants to “assemble multiple components into a unit, which is made by placing a peg in a hole (dominant hand), placing a washer over the peg (nondominant hand), then a small cylindrical collar (dominant hand), followed by a second washer on top (non-dominant hand)” (Wilkes et al., 2023). This engages fine motor function as well as sequencing skills and bimanual coordination. A higher value for Purdue Pegboard Assembly can be interpreted as better performance on this task.

#### 1.2.4. Shopping List Memory Task

The Shopping List Memory Task was administered on a computer. Words of 30 items resembling those commonly seen on a shopping list were presented to the participant one at a time for 14 seconds each with a 0.5 second fixation cross in between words. Memory for the items was assessed approximately 20 minutes later via recognition discrimination for the original list via percentage of items answered correctly. A higher value indicated better performance on this task. Outliers with values greater than 3 standard deviations from the mean were excluded from these task-based analyses (n=2).

### 1.3. Hormone quantification

We followed the methodology described in our recent work^54^ for hormonal analyses, and to ensure clarity and replicability, we have shared those methods directly here. Participants were instructed to abstain from alcohol consumption for 24 hours and to avoid eating or drinking for 3 hours prior to the first study session to minimize external influences on hormone levels. Saliva samples were collected exclusively in the morning and afternoon. Participants were also screened for oral disease or injury, use of substances such as nicotine or caffeine, and prescription medications that may impact the saliva pH and compromise samples. Participants rinsed their mouths with water 10 minutes before providing a saliva sample to remove any residue.

Samples were then collected in pre-labeled cryovials provided by Salimetrics (https://salimetrics.com/saliva-collection-training-videos/) using the passive drool technique. Participants were asked to supply 1mL of saliva, after which samples were immediately stored in a -80° Celsius bio-freezer for stabilization. Assays were completed by Salimetrics to quantify 17β-estradiol and progesterone levels for each participant. The amount of saliva collected was sufficient to detect 17β-estradiol at a high sensitivity threshold of 0.1pg/mL, along with 5.0 pg/mL and 1.0 pg/mL thresholds for progesterone and testosterone, respectively.

### 1.4. Imaging acquisition

The imaging sections (section 1.4 through 1.6) utilize standardized text to ensure consistency and reproducibility in the research protocol. These standardized descriptions can also be found in other publications from our laboratory^55,66^ and align with current best practices in the field, providing a clear and detailed framework for the procedures undertaken.

Participants underwent structural and resting-state MRI using a Siemens Magnetom Verio 3.0 Tesla scanner and a 32-channel head coil. For structural MRI, we collected a high-resolution T1-weighted 3D magnetization prepared rapid gradient multi-echo (MPRAGE) scan (repetition time (TR) = 2400 ms; acquisition time = 7 minutes; voxel size = 0.8 mm^3^) and a high-resolution T2-weighted scan (TR = 3200 ms; acquisition time = 5.5 minutes; voxel size = 0.8 mm^3^), each with a multiband acceleration factor of 2. For resting-state imaging, we administered four blood -oxygen level dependent (BOLD) functional connectivity (fcMRI) scans with the following parameters: multiband factor of 8, 488 volumes, TR of 720 ms, and 2.5 mm^3^ voxels. Each fcMRI scan was 6 minutes in length for a total of 24 minutes of resting-state imaging, and scans were acquired with alternating phase encoding directions (i.e., two anterior to posterior scans and two posteriors to anterior scans). During the fcMRI scans, participants were asked to lie still with their eyes open while fixating on a central cross. In total, the acquisition of images took about 45 minutes, including a 1.5-minute localizer.

Scanning protocols were adapted from the multiband sequences developed by the Human Connectome Project (HCP)^67^ and the Center for Magnetic Resonance Research at the University of Minnesota to facilitate future data sharing and reproducibility.

### 1.5. Image processing

The images underwent several preprocessing steps to prepare them for further analysis. Initially, they were converted from DICOM to NIFTI format and organized following the Brain Imaging Data Structure (BIDS, version 1.6.0) using the bidskit docker container (version 2021.6.14, https://github.com/jmtyszka/bidskit). Afterward, a single volume was extracted from two oppositely coded BOLD images to estimate B0 field maps using the split tool from the FMRIB Software Library (FSL) package (Jenkinson et al., 2012). Subsequently, the anatomical and functional images were preprocessed using fMRIPrep (version 20.2.3; for detailed methods, see https://fmriprep.org/), which includes automated procedures to align the functional volume with the anatomical image, correct for motion, correct field map distortions, segment the anatomical image into distinct tissues (e.g., gray matter, white matter, cerebrospinal fluid), remove the skull from the anatomical image, normalize the data to a common space, align motion-corrected functional volumes with the normalized anatomical image, and apply spatial smoothing.

### 1.6. Functional analysis

Following preprocessing with fMRIPrep, the subsequent analyses were conducted using the CONN toolbox (version 21a)^68^. This involved additional processing to eliminate noise and artifacts and enhance data quality. Denoising in CONN typically comprises several stages, such as removing motion signals and regressing out confounding signals (e.g., signals from ventricles, white matter, and global signals). Motion information from fMRIPrep was utilized in CONN for this purpose. A 0.008-0.099 Hz bandpass filter was applied to eliminate high-frequency noise. The denoising step is crucial for enhancing the quality of FC data by minimizing artifacts and enhancing the ability to detect genuine FC patterns in the brain.

Resting-state FC analyses focused on regions of interest (ROIs) in both the hippocampus and the cerebellum (**Figure 1**). The hippocampal seed regions were loosely based on coordinates derived from Grady’s meta-analysis on hippocampal function^69^. These coordinates helped determine anatomical hippocampal boundaries and the original coordinates were shifted from the original values to accommodate a new center for 5mm spherical seeds in MNI space. That is, certain coordinates from Grady^69^ fell on the outer edges of the hippocampus and to cover 3-dimensional space within the boundaries of the hippocampus, we had to shift certain coordinates to account for a 5mm spherical seed. Cerebellar seeds were determined by mapping dorsal, ventral, rostral, caudal, medial, and lateral boundaries of the right cerebellum in MNI space. Please refer to **Supplementary Table 2** for coordinates for all our seeds. Following this step, we used the oro.nifti and spatstat packages in R Studio to create a non-overlapping grid in 3 dimensional space of 5mm spherical seeds covering the right cerebellum^70,71^. As our aim for this study was to cover the entirety of the right cerebellum and left hippocampus, seeds were created to cover the 3-dimensional anatomical regions in MNI space. For ease of interpretation, we used the automated anatomical atlas (AAL) in the label4MRI package in R to approximate anatomical regions for each seed ^72^ which is specified in **Supplementary Table 2**. Our seeds are visualized in a simpler, easier to view, rendering in **Figure 1**, and we have created a more in-depth representation in **Supplementary** Figure 1. The investigation was limited to the right cerebellar hemisphere and left hippocampus, to mitigate multiple comparisons and ensure examination of cross lateralized connections.

**Figure 1.**
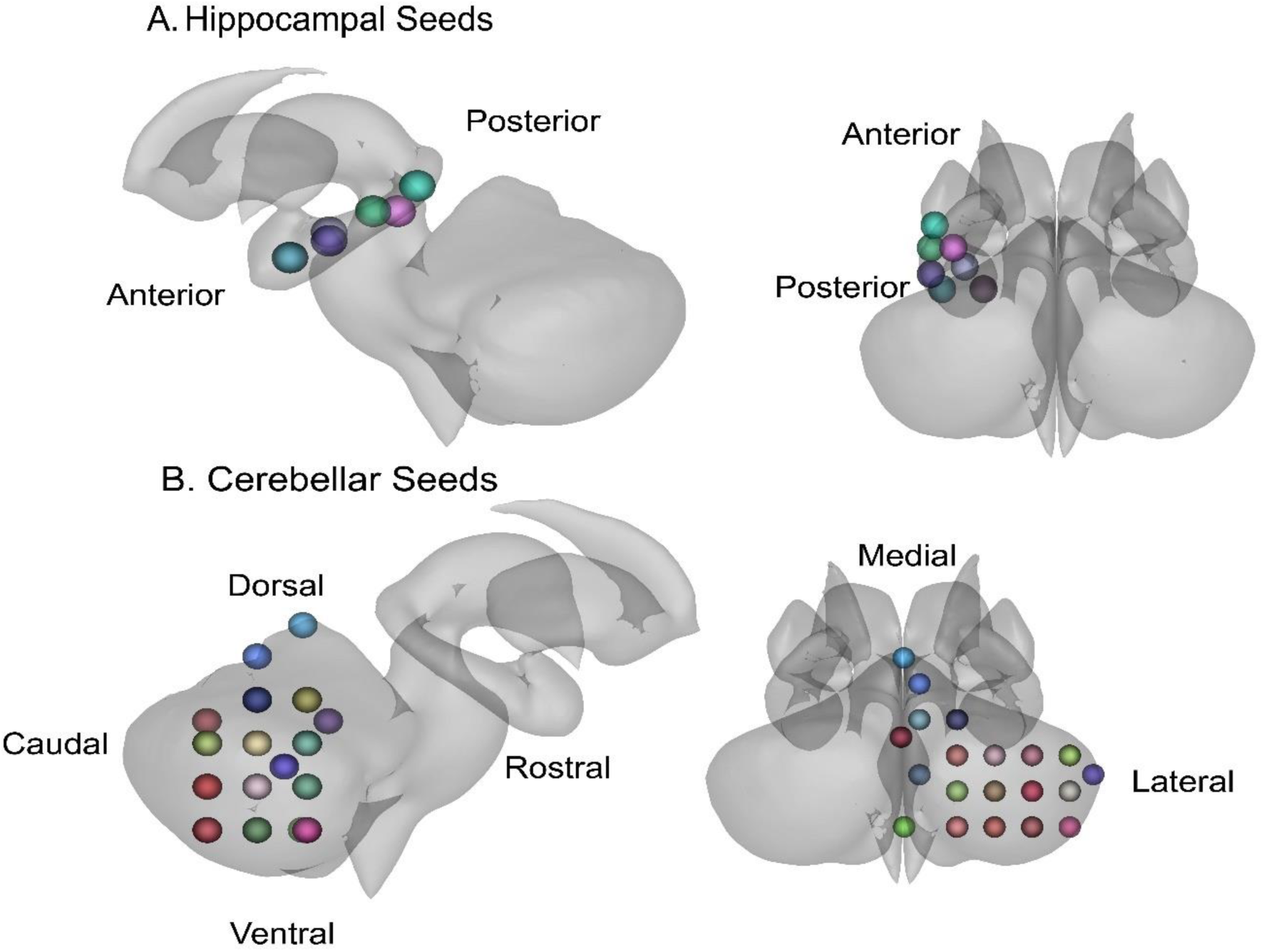
These figures illustrate several regions of interest (ROIs) used in this study. Seed colors are helpful for differentiating seeds, but do not hold additional meaning. **A**. Hippocampal seeds are displayed with directional indicators of the anterior and posterior long-axis. **B**. Cerebellar seeds are displayed with directional indicators for the right cerebellar hemisphere. The caudal region of the cerebellum is better visualized in a more detailed representation of the ROI seeds in **Supplementary Figure 1**.

CB-HP FC was examined via region of interest (ROI)-to-ROI correlations using CONN toolbox at the group level. Relationships between age and CB-HP FC were evaluated via correlations in the whole sample. Sex differences in CB-HP FC were investigated via an analysis of covariance (ANCOVA), controlling for age. Group-level correlations were also performed to examine CB-HP connectivity patterns in relation to hormone levels (i.e., 17-β estradiol, progesterone, respectively) in the whole sample, controlling for age. To determine brain-behavior performance relationships, the variables Symbol Span, Stroop, Purdue Pegboard Assembly, Shopping List Memory task, Sequence Learning, and Letter-Number Sequencing were separately correlated with CB-HP FC while controlling for age. We employed standard settings for cluster-based inferences using parametric statistics based on random field theory. We used an initial voxel threshold at p<.001 along with a cluster threshold set at p < .05, with a false discovery rate (FDR) correction. To determine effect size specifically with Cohen’s d, we utilized the equation ((2* t-value)/√degrees of freedom)^73^ in Excel. T-values and degrees of freedom were extracted from analytic results in CONN toolbox.

### 1.7. Statistical analyses

For statistical analyses of the age, behavior, and hormone level results, we used R (v2024.04.2.764, R Core Team, 2021). We first sought to ascertain relationships between age, education, and cognitive performance (i.e., Symbol Span, Stroop, Purdue Pegboard Assembly, Shopping List Memory task, Sequence Learning, and Letter-Number Sequencing) while controlling for education level via linear regressions. To explore differences in hormone levels (i.e., 17-β estradiol, progesterone) with age, we conducted linear regressions.

## Results

For the following results, please refer to **Figure 1** as a reference for directionality regarding both the hippocampus and cerebellum.

### Age is Associated with CB-HP Connectivity

Age was significantly correlated with CB-HP connectivity (**Figure 2, Supplementary Table 3**). Increased age correlated significantly with lower CB-HP FC; specifically encompassing the majority of the hippocampus on the anterior to posterior long axis and dorsal, medial, and caudal regions of the cerebellum (**Figure 2**). However, there were also findings in the opposite direction. Higher CB-HP FC correlated with increased age between the left posterior hippocampus and right dorsal cerebellum only.

**Figure 2.**
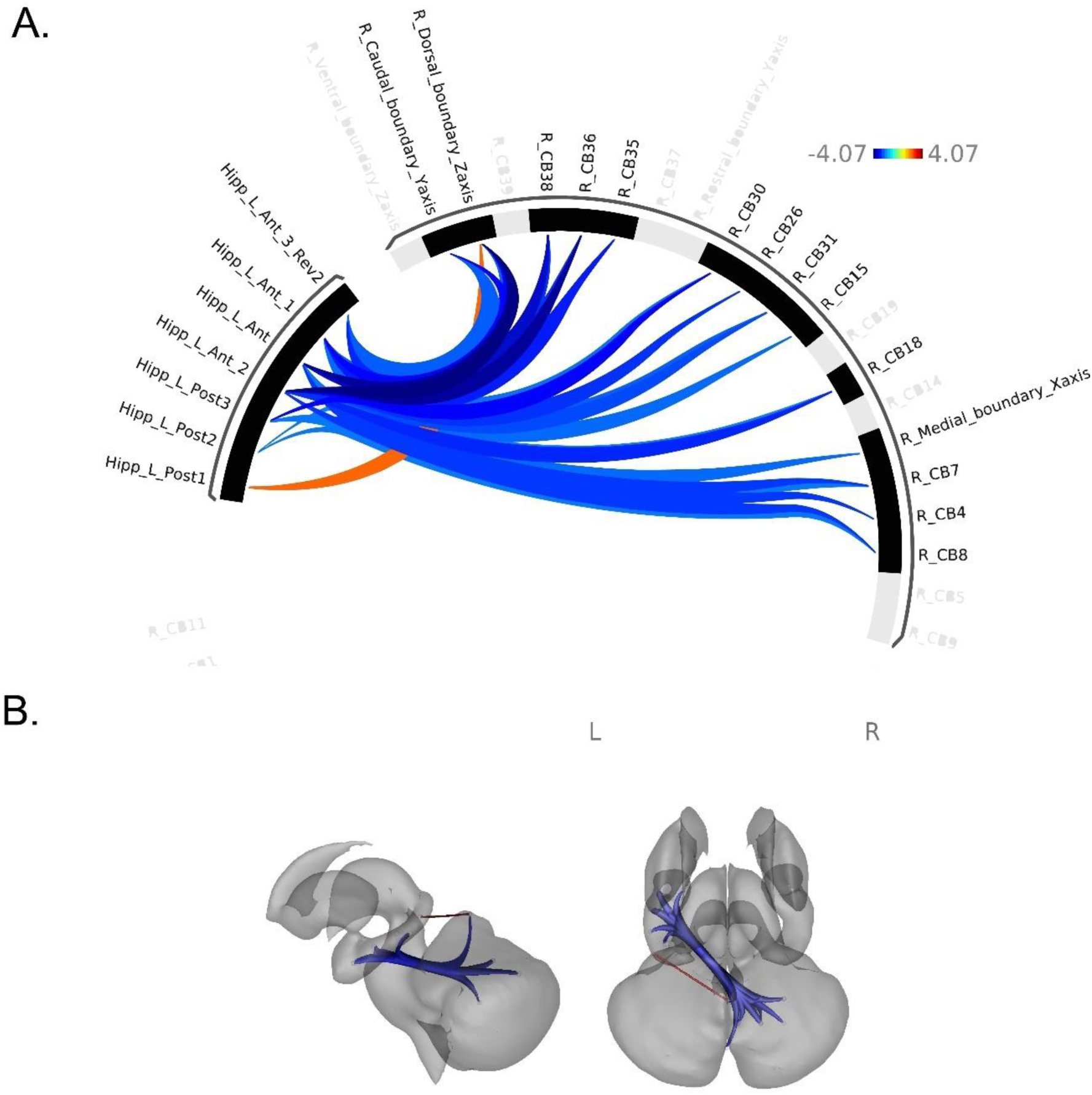
Patterns of cortical functional connectivity (FC) between ROIs. **A**. ROIs are shown in an FC ring where blue represents lower FC between ROIs with increased age; while orange-red displays greater FC between ROIs with increased age. **B**. ROIs are shown on a subcortical model where blue represents negative FC relationships between ROIs with increased age, while red displays positive FC relationships between ROIs with increased age. Colors in figure B do not correspond to the key in figure A and only indicate positive versus negative relationships in blue and red as described.

### CB-HP Connectivity and Age Associations with Hormone Levels

Sex differences in CB-HP FC were explored via group-level ROI-to-ROI contrasts in males versus females when controlling for age. We did not find significant sex differences in CB-HP FC (pFDR > 0.05). While the primary aim of our study was not intended to examine intracerebellar or intrahippocampal FC as we were interested in the CB-HP circuit, we did find greater intracerebellar FC in males as compared to females when controlling for age (pFDR < 0.05, **Supplementary** Figure 2).

We discovered several robust associations between hormones and CB-HP connectivity across all participants (male and female). Higher 17β-estradiol levels correlated significantly with higher CB-HP FC; more specifically, connectivity along the majority of the hippocampus on the anterior to posterior long axis and both medial and dorsal regions of the cerebellum were associated with higher levels of 17β-estradiol (**Figure 3, Supplementary Table 4**). However, there was also one finding in the opposite direction. Lower CB-HP FC correlated with greater 17β-estradiol levels between the left posterior hippocampus and a medial-dorsal region of the cerebellum (lobule VI) only. Regarding progesterone levels, we primarily found that higher progesterone levels were linked to lower CB-HP FC across the majority of both cerebellar and hippocampal ROIs, when controlling for age (**Figure 4, Supplementary Table 5**), the opposite of what we found with 17β-estradiol. We also demonstrated patterns of greater intracerebellar and intrahippocampal FC (respectively) with higher progesterone levels when controlling for age (**Supplementary** Figure 3).

**Figure 3.**
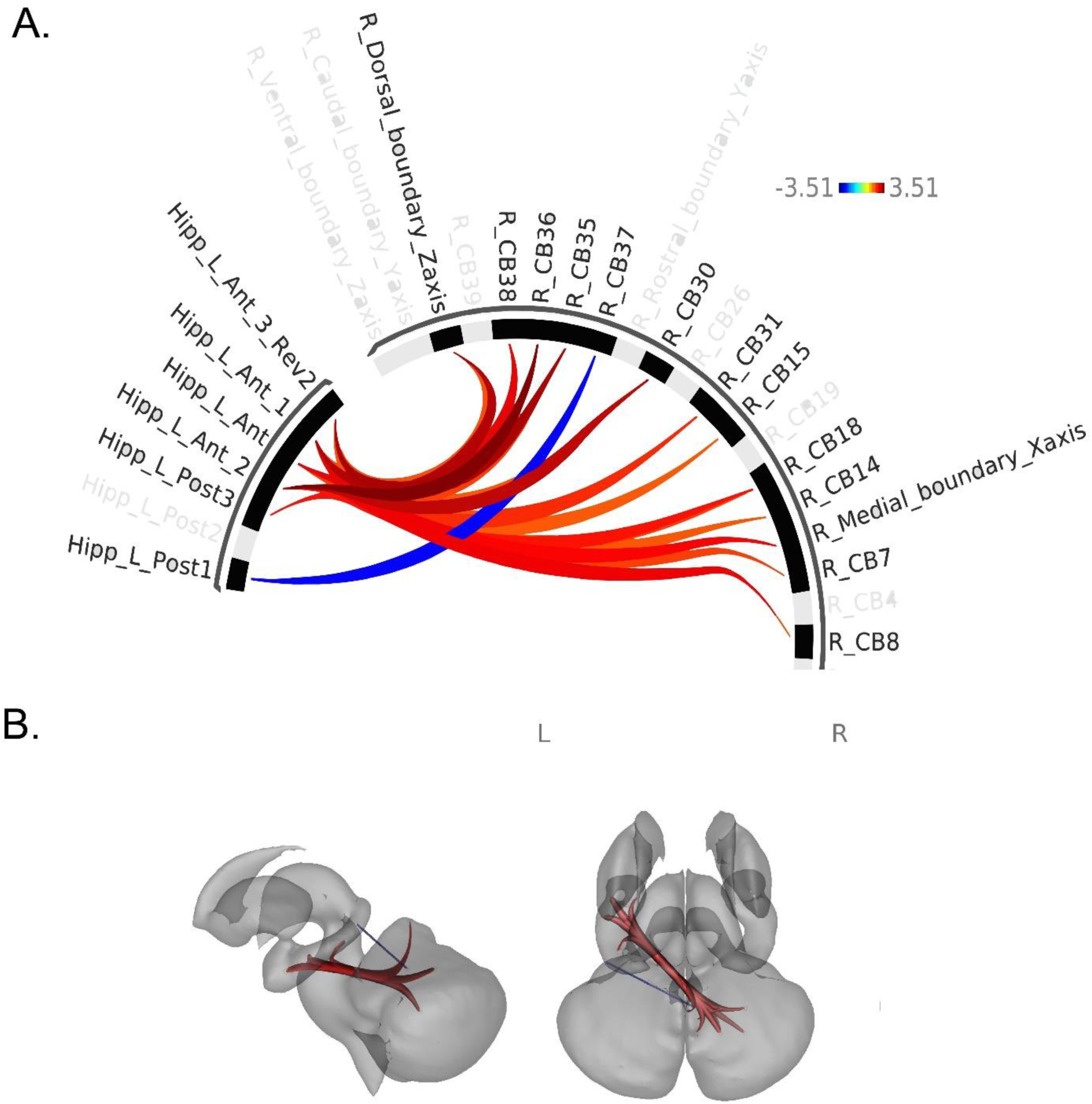
Associations between 17β-estradiol and CB-HP connectivity. **A**. ROIs are shown in an FC ring where orange-red displays greater FC between ROIs with higher levels of 17β-estradiol. Blue represents lower FC between ROIs with higher levels of 17β-estradiol. **B**. ROIs are shown on a subcortical model where red displays positive FC relationships, while blue represents negative FC relationships between ROIs with higher levels of 17β-estradiol. Colors in figure B do not correspond to the key in figure A and only indicate positive versus negative relationships in blue and red as described.

**Figure 4.**
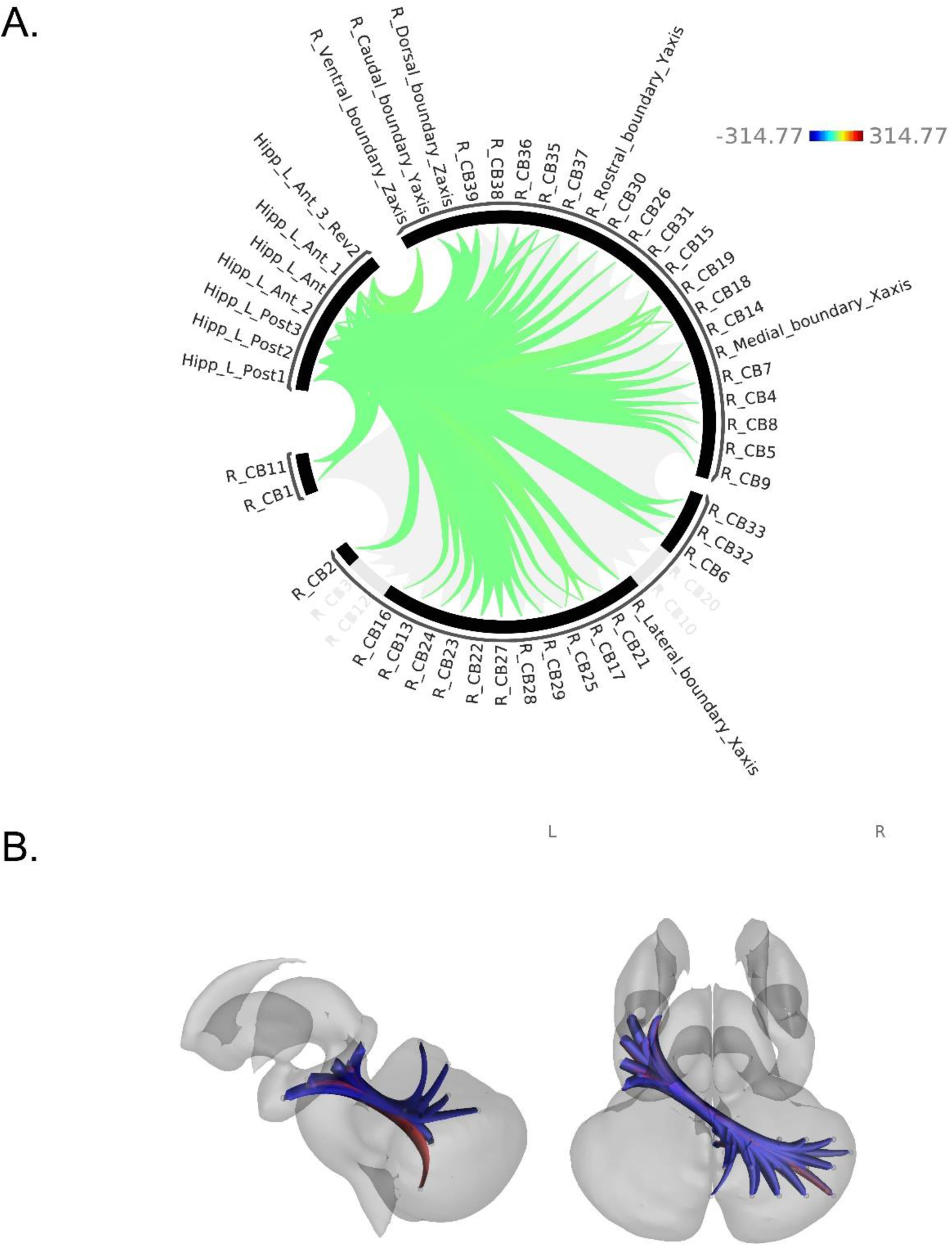
Associations between progesterone and CB-HP connectivity **A**. ROIs are shown in an FC ring where green displays lower FC between ROIs with higher levels of progesterone. **B**. ROIs are shown on a subcortical model where red displays positive FC relationships with higher levels of progesterone, while blue represents negative FC relationships with higher levels of progesterone. Colors in figure B do not correspond to the key in figure A and only indicate positive versus negative relationships in blue and red as described.

To assess for expected trends in our data, we evaluated linear relationships between hormone levels (i.e., 17β-estradiol and progesterone) and age. As expected, both 17β-estradiol (β = -0.0126 pFDR = 0.001) and progesterone (β = -1.7041 pFDR < 0.0001) displayed negative relationships with age.

### Age Associations with Behavioral Performance and CB-HP Connectivity

Regarding relationships between behavioral measures, age, and education level, linear regressions revealed significant negative relationships between age and Symbol Span, Shopping List Memory, and Letter-Number Sequencing, when controlling for education level (see **Supplementary Table 6** for statistics). That is, with increasing age, behavioral performance was significantly lower in Symbol Span, Shopping List Memory, and Letter-Number Sequencing. Relationships between Purdue Pegboard Assembly, Stroop Effect, Sequence Learning and age have been previously reported in this population^56^. There were no main effects of education level across behavioral tasks (pFDR > 0.05, **Supplementary Table 6**). There were no significant associations between CB-HP FC and cognitive variables (i.e., Symbol Span, Stroop, Pegboard Assembly, Shopping List Memory, Sequence Learning, or Letter-Number Sequencing) when controlling for age, pFDR > 0.05.

## Discussion

This study explored the CB-HP circuit in the context of age, hormone levels and sex differences, and behavioral performance across the adult lifespan. We have mapped this novel and important circuit for the first time, in detail, in the adult and aging human brain. Given the importance of these regions individually for behavior and aging, this work provides a critical mapping of the CB-HP circuit that stands to serve as a foundation for future work in age-related neurodegenerative disease. Our results revealed CB-HP functional associations with cross-sectional age, such that as age is higher, connectivity is lower. We also found associations with hormone levels, though the directionality of these relationships differs between 17β-estradiol and progesterone. Our study provides unique insights into the CB-HP circuit with respect to hormonal and behavioral differences across a wide age range of healthy middle aged and older adults.

### Hormone Level Associations with Age and CB-HP Connectivity

As expected, both 17-β estradiol and progesterone exhibited lower levels with increased age across participants. In women, 17-β estradiol levels fluctuate cyclically during the reproductive stage and drop significantly around perimenopause^46,74,75^. The trajectory of estradiol has shown a much lower rate of decline in aging men^75,76^. Progesterone levels tend to have the steepest decline around post-menopause in women and also display a lower rate of decline in aging men^75^. Specific sex differences in hormone levels (17-β estradiol and progesterone) were previously investigated by age group (early middle-age, late middle-age, and older adults) within the same sample as this study^57^ and aligned with the aforementioned literature. While not surprising, this provides an important confidence check with respect to the assays and results used here.

Consistent with our hypothesis and the literature on estradiol levels related to FC^28–31^, we found greater CB-HP connectivity with higher estradiol levels across participants when controlling for age. Previously, a review by Peper et. al (2011) noted that higher levels of both exogenous and endogenous estradiol were associated with increased FC within the cortex and between the cortex and subcortex^28^. In women, there is also evidence of greater coherence of functional networks with higher levels of 17-β estradiol^29,30^. However, in a dense sampling of a single reproductive aged woman, higher estradiol was associated with reductions in cerebellar connectivity to other networks^34^. We specifically found greater FC between the majority of the hippocampus along the anterior-posterior long axis and both medial and dorsal regions of the cerebellum. Based on 7-network parcellation of the cerebellum by Yeo and colleagues (2011), the medial-dorsal regions of the cerebellum that showed greater FC to the hippocampus at rest in this study, display overlap with somatomotor, limbic, and ventral attention networks^31^. This finding is consistent with the hippocampus’ key role in limbic network function. While the majority of significant associations demonstrated lower connectivity with higher levels of 17β-estradiol, we did see a positive correlation such that connectivity between the left posterior hippocampus and a medial-dorsal region of the cerebellum were higher instead. This cerebellar region overlapped with the limbic network^31^, which was unexpected. However, the relationship we found between higher 17β-estradiol and greater FC in part of the limbic network could have affective implications that were not explored in this study.

Overall, our results showing lower CB-HP FC with higher levels of progesterone add evidence to the literature suggesting lower cerebello-cortical FC with higher endogenous progesterone across aging men and women. Further, we found greater intracerebellar FC with higher levels of progesterone. However, direct comparisons between our results and findings to date are limited to young women in their reproductive stage^30^ as well as dense samplings (n=1) of that same population^29,32–35^. Dense sampling revealed reduced network coherence with higher levels of progesterone across 7 out of 9 functional networks examined ^29^ which aligns with our findings of reduced CB-HP FC. However, another dense sampling in a reproductive stage woman displayed increased FC between the hippocampus, dorsolateral prefrontal cortex, and sensorimotor cortex with increased progesterone levels ^32^. Compounding the mixed results in the literature, a small group of reproductive aged women displayed both positive and negative correlations between intracortical FC and progesterone levels^30^. Thus, there are not clear patterns in the literature to draw direct conclusions about our results. Broadly, our results suggest that progesterone modulates the CB-HP circuit and intracerebellar FC in both sexes and across the adult lifespan. While our behavioral results did not support our hypothesis that the FC relationships seen here are related to preclinical cognitive function, our results may illuminate important indicators of brain function in aging and the impact of the hormonal environment on brain function whether or not there are sex differences. For instance, progesterone has been described as an antagonist for estradiol’s proliferative impact on dendritic spines^77^, suggesting that ratios between estradiol and progesterone could provide valuable insight on brain network organization, characterization, and ultimate associations with behavioral functioning.

Our results have particular implications for aging women given their significant declines in sex steroid hormones with aging^78^. Diminished estradiol has also been linked to greater AD risk and pathogenesis in the aging brain^79–83^. Our findings of lower CB-HP FC with higher progesterone levels and greater CB-HP FC with higher 17β-estradiol levels suggest that these hormones modulate brain function in a circuit that appears to have relevance in aging and AD risk. Future research could benefit from targeted investigation of this circuit in the context of menopausal transition in women and in age-related neurodegenerative disease.

### Age is Associated with CB-HP Connectivity

In aging and Alzheimer’s Disease (AD), hippocampal structure and function are notably impacted^84,85^. Recently, cerebellar integrity has also been implicated in aging and AD via neuroimaging studies and amyloid-ß presence in familial cases^86–91^. While the cerebellum may not be a primary driver of pathology in aging, evidence suggests that it plays a significant role in symptomatology through its cognitive and behavioral interactions, particularly noted in cerebellar cognitive affective syndrome^92^ A recent review by Bernard (2024) suggests that this cerebellar scaffolding in aging supports optimal cortical function, but as AD progresses, its effectiveness diminishes, pointing to the need for more research on cerebellar-hippocampal interactions^93^. Taken together, the literature implies that the CB-HP circuit plays a role aging and AD.

Consistent with our hypothesis, greater age was associated with lower CB-HP FC across most of the nodes we investigated here. However, we did demonstrate one instance where CB-HP FC was higher with older age when looking at the left posterior hippocampus and right dorsal cerebellum only. A comparison of young versus older adults similarly exhibited lower FC between several cerebellar regions and both the hippocampus and parahippocampal gyrus in older adults^3^. In healthy aging adults, Uwisengeyimana and colleagues found alterations in cerebellar to hippocampal FC across five age groups that started at 40 years of age and ended at 90 years which is somewhat consistent with the patterns of both lower and higher CB-HP FC with increased age seen here ^94^. To date, human studies of the CB-HP FC have largely been limited to visuospatial task-based examinations in young adults^20,26,27^. Our approach is the first to evaluate this circuit at rest and across a healthy aging population.

Notably, studies with broader explorations that included the hippocampus and cerebellum found FC relationships when comparing healthy older adult populations to those with significant neuropathology (AD) and/or cognitive disfunction (mild cognitive impairment; MCI)^95–98^. While these studies did not specifically target the CB-HP circuit, those specifically comparing AD or amnestic MCI patients to healthy controls found lower FC between the hippocampus and cerebellum^97,98^, consistent with our hypothesis that CB-HP FC declines with aging and is impacted by neurodegenerative pathology. In conceptualizing the literature here comparing different stages of adulthood, MCI, or AD^95–98^, we speculate a quadratic relationship with CB-HP as its related to neuropathology, cognitive function, and functional abilities. Specifically, lower CB-HP FC in younger adulthood, increased compensatory FC^99^ in middle-age or at the MCI stage, and declines in CB-HP in the context of normative aging pathology or progressed neuropathology (e.g., AD).

### Behavioral Associations with Age, Sex differences, Hormones, and CB-HP Connectivity

Regarding CB-HP FC associations with behavioral measures, we did not show significant associations across tasks. Task-based cerebellar functional activation has been seen in attention, working memory, language, executive function, and spatial abilities^100,101^. Task-based hippocampal functional activation has shown the strongest associations with encoding, memory retrieval, and spatial processing ^1,102,103^. However, our results were not consistent with studies implicating a CB-HP circuit in spatial, cognitive-motor, and episodic memory tasks^17,20,98^. This may be attributed to our cognitive measures as they were initially intended to examine cerebellar function, and tasks that are easily translatable to those given in clinical cognitive assessments. Measures that are more sensitive to spatio-temporal relationships, visuomotor integration, and more complex measures of episodic memory may be required to appropriately assess the CB-HP circuit as it relates to behavior. Another limitation was that behavioral tasks were administered outside of the scanner. It is possible that cognitive tasks in the domains we examined would engage the CB-HP circuit if they were assessed with fMRI. Future work examining the CB-HP circuit using a task-based paradigm is warranted and may clarify which tasks are most sensitive to changes in this circuit with aging.

## Conclusion

Our results characterized relationships between age, the CB-HP circuit, behavioral performance, sex differences, and sex hormone levels across the adult lifespan. We found largely lower CB-HP FC with increased age and poorer behavioral performance with increased age. Both estradiol and progesterone appear to modulate CB-HP FC, with higher estradiol largely associated with greater FC and progesterone largely associated with lower FC. Given the notable changes in sex steroid hormone levels with menopause in aging females, this is of particular interest and potential importance. Most notable are possible implications for this circuit in age-related neurodegenerative disease such as Alzheimer’s Disease, and sex differences therein. These results replicate and extend findings in the literature supporting a role for the CB-HP circuit in understanding advanced age. Future research can utilize these findings as a foundation for clinical and translational applications, given the robust baseline established by the wide age range and varied set of tasks used in this study.

## Acknowledgment

This work was supported by R01AG065010 to J.A.B. This work was further supported by the Texas Virtual Data Library (ViDaL), a high-performance cluster, funded by the Texas A&M University Research Development fund. In this cluster the imaging analyses for the current work were carried out using the resources provided by the Texas A&M High Performance Research Computing organization.

## Notes

### Competing Interest Statement

The authors have declared no competing interest.

