## Supplement for "The Human Cerebello-Hippocampal Circuit Across Adulthood"

**Supplementary Table 1**. Demographic means for the full sample are listed below with standard deviations in parentheses.

| Demographics | |
|---|---|
| Age (years) | 57(13.31) |
| Sex F(M) | 74(64) |
| Education (years) | 16.7 (2.3) |
| African American (%) | 2% |
| Asian (%) | 7% |
| Caucasian (%) | 86% |
| Multiracial (%) | 4% |
| Native American (%) | 1% |

**Supplementary Table 2.** Cerebellar and Hippocampal Coordinates for 5mm Spherical Seeds

| Seed Label | X | Y | Z | Region (AAL) |
|---|---|---|---|---|
| R_Medial boundary_Xaxis | 5.00 | -59.20 | -36.50 | Vermis IX |
| R_Lateral_boundary_Xaxis | 51.30 | -59.10 | -36.50 | Crus I |
| R_Caudal_boundary_Yaxis | 0.00 | -74.70 | -26.00 | Vermis VII |
| R_Rostral_boundary_Yaxis | 0.20 | -50.30 | -26.00 | Vermis X |
| R_Dorsal_boundary_Zaxis | 0.80 | -55.40 | -3.90 | Vermis IV-V |
| R_Ventral_boundary_Zaxis | 1.00 | -55.50 | -51.10 | Lobule IX |
| R_CB1 | 15.00 | -74.60 | -51.10 | Lobule VIII |
| R_CB2 | 25.00 | -74.60 | -51.10 | Lobule VIIb |
| R_CB3 | 35.00 | -74.60 | -51.10 | Lobule VIIb |
| R_CB4 | 15.00 | -64.60 | -51.10 | Lobule VIII |
| R_CB5 | 25.00 | -64.60 | -51.10 | Lobule VIII |
| R_CB6 | 35.00 | -64.60 | -51.10 | Lobule VIII |
| R_CB7 | 15.00 | -54.60 | -51.10 | Lobule IX |
| R_CB8 | 25.00 | -54.60 | -51.10 | Lobule VIII |
| R_CB9 | 35.00 | -54.60 | -51.10 | Lobule VIII |
| R_CB10 | 45.00 | -54.60 | -51.10 | Lobule VIIb |
| R_CB11 | 15.00 | -74.60 | -41.10 | Lobule VIIb |
| R_CB12 | 25.00 | -74.60 | -41.10 | Crus II |
| R_CB13 | 35.00 | -74.60 | -41.10 | Crus II |
| R_CB14 | 15.00 | -64.60 | -41.10 | Lobule VIII |
| R_CB15 | 25.00 | -64.60 | -41.10 | Lobule VIII |
| R_CB16 | 35.00 | -64.60 | -41.10 | Crus II |
| R_CB17 | 45.00 | -64.60 | -41.10 | Crus II |
| R_CB18 | 15.00 | -54.60 | -41.10 | Lobule VIII |
| R_CB19 | 25.00 | -54.60 | -41.10 | Lobule VIII |
| R_CB20 | 35.00 | -54.60 | -41.10 | Crus I |
| R_CB21 | 45.00 | -54.60 | -41.10 | Crus II |

| | | | | |
|---|---|---|---|---|
| R_CB22 | 15.00 | -74.60 | -31.10 | Crus I |
| R_CB23 | 25.00 | -74.60 | -31.10 | Crus I |
| R_CB24 | 35.00 | -74.60 | -31.10 | Crus I |
| R_CB25 | 45.00 | -74.60 | -31.10 | Crus I |
| R_CB26 | 15.00 | -64.60 | -31.10 | Lobule VI |
| R_CB27 | 25.00 | -64.60 | -31.10 | Lobule VI |
| R_CB28 | 35.00 | -64.60 | -31.10 | Crus I |
| R_CB29 | 45.00 | -64.60 | -31.10 | Crus I |
| R_CB30 | 15.00 | -54.60 | -31.10 | Lobule VI |
| R_CB31 | 25.00 | -54.60 | -31.10 | Lobule VI |
| R_CB32 | 35.00 | -54.60 | -31.10 | Crus I |
| R_CB33 | 45.00 | -54.60 | -31.10 | Crus I |
| R_CB34 | 15.00 | -74.60 | -21.10 | Lobule VI |
| R_CB35 | 5.00 | -64.60 | -21.10 | Vermis VI |
| R_CB36 | 15.00 | -64.60 | -21.10 | Lobule VI |
| R_CB37 | 5.00 | -54.60 | -21.10 | Vermis IV-V |
| R_CB38 | 15.00 | -54.60 | -21.10 | Lobule VI |
| R_CB39 | 5.00 | -64.60 | -11.10 | Vermis VI |
| Hipp_L_Ant | -30.00 | -10.00 | -22.00 | Hippocampus |
| Hipp_L_Ant_1 | -24.00 | -18.00 | -16.00 | Hippocampus |
| Hipp_L_Ant_2 | -33.00 | -18.00 | -18.00 | Hippocampus |
| Hipp_L_Ant_3_Rev2 | -19.00 | -10.00 | -22.00 | Parahippocampal gyrus |
| Hipp_L_Post1 | -32.00 | -36.00 | -5.00 | Hippocampus |
| Hipp_L_Post2 | -33.00 | -27.00 | -11.00 | Hippocampus |
| Hipp_L_Post3 | -27.00 | -32.00 | -11.00 | Parahippocampal gyrus |

**Supplementary Figure 1.** Cerebellar Seeds Rendered in marsbar.

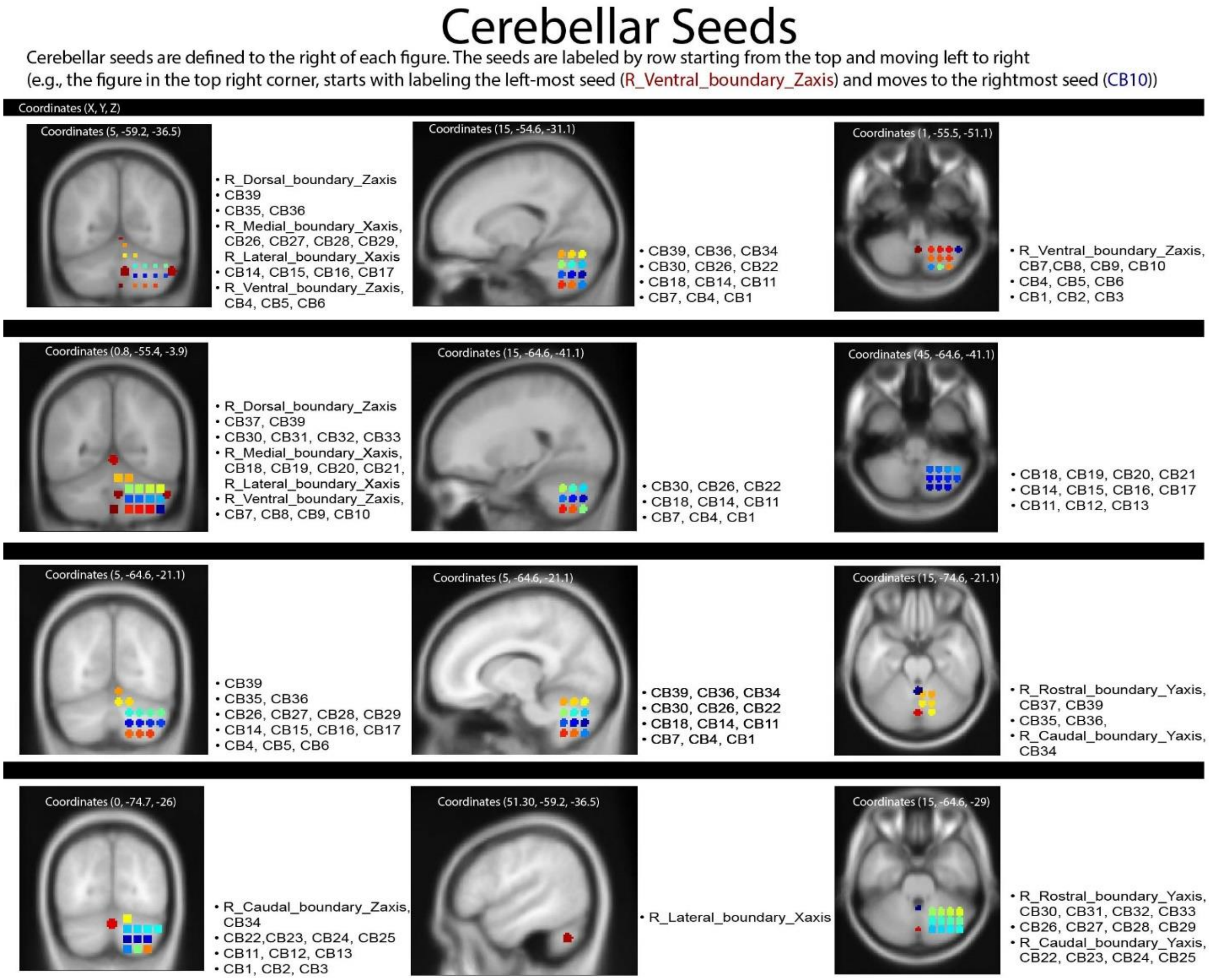


**Supplementary Table 3.** Cerebellar-hippocampal regions showing significant correlations in cortical FC with age.

| FC Seed Regions | | T(136) | pFDR | Cohen's d |
|---|---|---|---|---|
| R_Dorsal_boundary_Zaxis | Hipp_L_Ant_2 | -4.07 | 0.002382 | -0.70 |
| Hipp_L_Ant_2 | R_Dorsal_boundary_Zaxis | -4.07 | 0.002382 | -0.70 |
| Hipp_L_Ant_2 | R_CB36 | -3.84 | 0.002382 | -0.66 |
| R_Dorsal_boundary_Zaxis | Hipp_L_Ant | -3.82 | 0.002382 | -0.66 |
| R_CB36 | Hipp_L_Ant_2 | -3.84 | 0.002382 | -0.66 |
| Hipp_L_Ant | R_Dorsal_boundary_Zaxis | -3.82 | 0.002382 | -0.66 |
| Hipp_L_Ant | R_CB38 | -3.49 | 0.005873 | -0.60 |
| Hipp_L_Ant | R_CB36 | -3.33 | 0.005873 | -0.57 |
| R_CB38 | Hipp_L_Ant | -3.49 | 0.00805 | -0.60 |
| R_CB36 | Hipp_L_Ant | -3.33 | 0.00805 | -0.57 |

| | | | | |
|---|---|---|---|---|
| Hipp_L_Ant_2 | R_CB30 | -3.07 | 0.013939 | -0.53 |
| R_Dorsal_boundary_Zaxis | Hipp_L_Ant_1 | -3.05 | 0.013939 | -0.52 |
| Hipp_L_Ant_2 | R_CB38 | -2.81 | 0.013939 | -0.48 |
| Hipp_L_Ant_2 | R_CB18 | -2.77 | 0.013939 | -0.48 |
| Hipp_L_Ant | R_CB4 | -2.64 | 0.014398 | -0.45 |
| Hipp_L_Ant | R_CB7 | -2.62 | 0.014398 | -0.45 |
| Hipp_L_Ant_2 | R_CB26 | -2.67 | 0.016437 | -0.46 |
| Hipp_L_Ant_2 | R_CB8 | -2.62 | 0.016437 | -0.45 |
| R_CB36 | Hipp_L_Ant_1 | -2.88 | 0.016437 | -0.49 |
| R_CB36 | Hipp_L_Ant_3_Rev2 | -2.76 | 0.016437 | -0.47 |
| R_CB36 | Hipp_L_Post3 | -2.75 | 0.016437 | -0.47 |
| Hipp_L_Ant_2 | R_CB31 | -2.47 | 0.016437 | -0.42 |
| Hipp_L_Post3 | R_Caudal_boundary_Yaxis | -3.00 | 0.016437 | -0.51 |
| Hipp_L_Post3 | R_CB35 | -2.87 | 0.016437 | -0.49 |
| R_Caudal_boundary_Yaxis | Hipp_L_Post3 | -3.00 | 0.016437 | -0.51 |
| Hipp_L_Post3 | R_CB36 | -2.75 | 0.016437 | -0.47 |
| R_CB38 | Hipp_L_Ant_2 | -2.81 | 0.016437 | -0.48 |
| Hipp_L_Ant_2 | R_CB15 | -2.22 | 0.016437 | -0.38 |
| R_CB35 | Hipp_L_Post3 | -2.87 | 0.016437 | -0.49 |
| R_CB35 | Hipp_L_Ant_3_Rev2 | -2.81 | 0.016437 | -0.48 |
| R_CB35 | Hipp_L_Ant_1 | -2.71 | 0.016877 | -0.46 |
| Hipp_L_Ant_1 | R_Dorsal_boundary_Zaxis | -3.05 | 0.016877 | -0.52 |
| Hipp_L_Ant_1 | R_CB36 | -2.88 | 0.017632 | -0.49 |
| Hipp_L_Ant | R_Medial_boundary_Xaxis | -2.24 | 0.017632 | -0.38 |
| Hipp_L_Ant_1 | R_CB35 | -2.71 | 0.017632 | -0.46 |
| R_CB30 | Hipp_L_Ant_2 | -3.07 | 0.017632 | -0.53 |
| R_Dorsal_boundary_Zaxis | Hipp_L_Ant_3_Rev2 | -2.34 | 0.017632 | -0.40 |
| R_Dorsal_boundary_Zaxis | Hipp_L_Post1 | 2.25 | 0.017632 | 0.39 |
| Hipp_L_Ant | R_CB18 | -2.04 | 0.017632 | -0.35 |
| Hipp_L_Post3 | R_CB38 | -2.48 | 0.017632 | -0.43 |
| R_CB18 | Hipp_L_Ant_2 | -2.77 | 0.024152 | -0.48 |
| Hipp_L_Ant_3_Rev2 | R_CB35 | -2.81 | 0.024152 | -0.48 |
| Hipp_L_Ant_3_Rev2 | R_CB36 | -2.76 | 0.024152 | -0.47 |
| R_CB38 | Hipp_L_Post3 | -2.48 | 0.024152 | -0.43 |
| Hipp_L_Ant_3_Rev2 | R_Dorsal_boundary_Zaxis | -2.34 | 0.031141 | -0.40 |
| R_Caudal_boundary_Yaxis | Hipp_L_Ant_1 | -2.34 | 0.031141 | -0.40 |
| R_Caudal_boundary_Yaxis | Hipp_L_Ant_3_Rev2 | -2.25 | 0.031141 | -0.39 |
| Hipp_L_Ant_3_Rev2 | R_Caudal_boundary_Yaxis | -2.25 | 0.031141 | -0.39 |
| Hipp_L_Ant_1 | R_Caudal_boundary_Yaxis | -2.34 | 0.034154 | -0.40 |
| Hipp_L_Ant_1 | R_CB31 | -2.28 | 0.034154 | -0.39 |
| Hipp_L_Ant_1 | R_CB30 | -2.16 | 0.034154 | -0.37 |
| R_CB31 | Hipp_L_Ant_2 | -2.47 | 0.034154 | -0.42 |
| R_CB31 | Hipp_L_Ant_1 | -2.28 | 0.034154 | -0.39 |
| Hipp_L_Post3 | R_CB8 | -2.17 | 0.034154 | -0.37 |
| Hipp_L_Post3 | R_CB7 | -2.12 | 0.034154 | -0.36 |

| | | | | |
|---|---|---|---|---|
| Hipp_L_Ant_1 | R_CB38 | -2.02 | 0.034154 | -0.35 |
| R_CB4 | Hipp_L_Ant | -2.64 | 0.034782 | -0.45 |
| R_CB8 | Hipp_L_Ant_2 | -2.62 | 0.034782 | -0.45 |
| R_CB8 | Hipp_L_Post3 | -2.17 | 0.038173 | -0.37 |
| R_CB8 | Hipp_L_Post2 | -2.09 | 0.038173 | -0.36 |
| R_CB7 | Hipp_L_Ant | -2.62 | 0.038173 | -0.45 |
| R_CB26 | Hipp_L_Ant_2 | -2.67 | 0.038173 | -0.46 |
| R_CB7 | Hipp_L_Post3 | -2.12 | 0.039912 | -0.36 |
| R_CB18 | Hipp_L_Ant | -2.04 | 0.039912 | -0.35 |
| R_CB38 | Hipp_L_Ant_1 | -2.02 | 0.042022 | -0.35 |
| R_CB38 | Hipp_L_Post2 | -1.99 | 0.042022 | -0.34 |
| R_CB30 | Hipp_L_Ant_1 | -2.16 | 0.045879 | -0.37 |
| R_CB15 | Hipp_L_Ant_2 | -2.22 | 0.045879 | -0.38 |
| R_Medial_boundary_Xaxis | Hipp_L_Ant | -2.24 | 0.046119 | -0.38 |
| Hipp_L_Post2 | R_CB8 | -2.09 | 0.046119 | -0.36 |
| Hipp_L_Post2 | R_CB38 | -1.99 | 0.048173 | -0.34 |
| Hipp_L_Post1 | R_Dorsal_boundary_Zaxis | 2.25 | 0.048173 | 0.39 |
| R_Dorsal_boundary_Zaxis | Hipp_L_Ant_2 | -4.07 | 0.002382 | -0.70 |
| Hipp_L_Ant_2 | R_Dorsal_boundary_Zaxis | -4.07 | 0.002382 | -0.70 |
| Hipp_L_Ant_2 | R_CB36 | -3.84 | 0.002382 | -0.66 |
| R_Dorsal_boundary_Zaxis | Hipp_L_Ant | -3.82 | 0.002382 | -0.66 |
| R_CB36 | Hipp_L_Ant_2 | -3.84 | 0.002382 | -0.66 |
| Hipp_L_Ant | R_Dorsal_boundary_Zaxis | -3.82 | 0.002382 | -0.66 |
| Hipp_L_Ant | R_CB38 | -3.49 | 0.005873 | -0.60 |
| Hipp_L_Ant | R_CB36 | -3.33 | 0.005873 | -0.57 |
| R_CB38 | Hipp_L_Ant | -3.49 | 0.00805 | -0.60 |
| R_CB36 | Hipp_L_Ant | -3.33 | 0.00805 | -0.57 |
| Hipp_L_Ant_2 | R_CB30 | -3.07 | 0.013939 | -0.53 |
| R_Dorsal_boundary_Zaxis | Hipp_L_Ant_1 | -3.05 | 0.013939 | -0.52 |
| Hipp_L_Ant_2 | R_CB38 | -2.81 | 0.013939 | -0.48 |
| Hipp_L_Ant_2 | R_CB18 | -2.77 | 0.013939 | -0.48 |
| Hipp_L_Ant | R_CB4 | -2.64 | 0.014398 | -0.45 |
| Hipp_L_Ant | R_CB7 | -2.62 | 0.014398 | -0.45 |
| Hipp_L_Ant_2 | R_CB26 | -2.67 | 0.016437 | -0.46 |
| Hipp_L_Ant_2 | R_CB8 | -2.62 | 0.016437 | -0.45 |
| R_CB36 | Hipp_L_Ant_1 | -2.88 | 0.016437 | -0.49 |
| R_CB36 | Hipp_L_Ant_3_Rev2 | -2.76 | 0.016437 | -0.47 |
| R_CB36 | Hipp_L_Post3 | -2.75 | 0.016437 | -0.47 |
| Hipp_L_Ant_2 | R_CB31 | -2.47 | 0.016437 | -0.42 |
| Hipp_L_Post3 | R_Caudal_boundary_Yaxis | -3.00 | 0.016437 | -0.51 |
| Hipp_L_Post3 | R_CB35 | -2.87 | 0.016437 | -0.49 |
| R_Caudal_boundary_Yaxis | Hipp_L_Post3 | -3.00 | 0.016437 | -0.51 |
| Hipp_L_Post3 | R_CB36 | -2.75 | 0.016437 | -0.47 |
| R_CB38 | Hipp_L_Ant_2 | -2.81 | 0.016437 | -0.48 |
| Hipp_L_Ant_2 | R_CB15 | -2.22 | 0.016437 | -0.38 |

| | | | | |
|---|---|---|---|---|
| R_CB35 | Hipp_L_Post3 | -2.87 | 0.016437 | -0.49 |
| R_CB35 | Hipp_L_Ant_3_Rev2 | -2.81 | 0.016437 | -0.48 |
| R_CB35 | Hipp_L_Ant_1 | -2.71 | 0.016877 | -0.46 |
| Hipp_L_Ant_1 | R_Dorsal_boundary_Zaxis | -3.05 | 0.016877 | -0.52 |
| Hipp_L_Ant_1 | R_CB36 | -2.88 | 0.017632 | -0.49 |
| Hipp_L_Ant | R_Medial_boundary_Xaxis | -2.24 | 0.017632 | -0.38 |
| Hipp_L_Ant_1 | R_CB35 | -2.71 | 0.017632 | -0.46 |
| R_CB30 | Hipp_L_Ant_2 | -3.07 | 0.017632 | -0.53 |
| R_Dorsal_boundary_Zaxis | Hipp_L_Ant_3_Rev2 | -2.34 | 0.017632 | -0.40 |
| R_Dorsal_boundary_Zaxis | Hipp_L_Post1 | 2.25 | 0.017632 | 0.39 |
| Hipp_L_Ant | R_CB18 | -2.04 | 0.017632 | -0.35 |
| Hipp_L_Post3 | R_CB38 | -2.48 | 0.017632 | -0.43 |
| R_CB18 | Hipp_L_Ant_2 | -2.77 | 0.024152 | -0.48 |
| Hipp_L_Ant_3_Rev2 | R_CB35 | -2.81 | 0.024152 | -0.48 |
| Hipp_L_Ant_3_Rev2 | R_CB36 | -2.76 | 0.024152 | -0.47 |
| R_CB38 | Hipp_L_Post3 | -2.48 | 0.024152 | -0.43 |
| Hipp_L_Ant_3_Rev2 | R_Dorsal_boundary_Zaxis | -2.34 | 0.031141 | -0.40 |
| R_Caudal_boundary_Yaxis | Hipp_L_Ant_1 | -2.34 | 0.031141 | -0.40 |
| R_Caudal_boundary_Yaxis | Hipp_L_Ant_3_Rev2 | -2.25 | 0.031141 | -0.39 |
| Hipp_L_Ant_3_Rev2 | R_Caudal_boundary_Yaxis | -2.25 | 0.031141 | -0.39 |
| Hipp_L_Ant_1 | R_Caudal_boundary_Yaxis | -2.34 | 0.034154 | -0.40 |
| Hipp_L_Ant_1 | R_CB31 | -2.28 | 0.034154 | -0.39 |
| Hipp_L_Ant_1 | R_CB30 | -2.16 | 0.034154 | -0.37 |
| R_CB31 | Hipp_L_Ant_2 | -2.47 | 0.034154 | -0.42 |
| R_CB31 | Hipp_L_Ant_1 | -2.28 | 0.034154 | -0.39 |
| Hipp_L_Post3 | R_CB8 | -2.17 | 0.034154 | -0.37 |
| Hipp_L_Post3 | R_CB7 | -2.12 | 0.034154 | -0.36 |
| Hipp_L_Ant_1 | R_CB38 | -2.02 | 0.034154 | -0.35 |
| R_CB4 | Hipp_L_Ant | -2.64 | 0.034782 | -0.45 |
| R_CB8 | Hipp_L_Ant_2 | -2.62 | 0.034782 | -0.45 |
| R_CB8 | Hipp_L_Post3 | -2.17 | 0.038173 | -0.37 |
| R_CB8 | Hipp_L_Post2 | -2.09 | 0.038173 | -0.36 |
| R_CB7 | Hipp_L_Ant | -2.62 | 0.038173 | -0.45 |
| R_CB26 | Hipp_L_Ant_2 | -2.67 | 0.038173 | -0.46 |
| R_CB7 | Hipp_L_Post3 | -2.12 | 0.039912 | -0.36 |
| R_CB18 | Hipp_L_Ant | -2.04 | 0.039912 | -0.35 |
| R_CB38 | Hipp_L_Ant_1 | -2.02 | 0.042022 | -0.35 |
| R_CB38 | Hipp_L_Post2 | -1.99 | 0.042022 | -0.34 |
| R_CB30 | Hipp_L_Ant_1 | -2.16 | 0.045879 | -0.37 |
| R_CB15 | Hipp_L_Ant_2 | -2.22 | 0.045879 | -0.38 |
| R_Medial_boundary_Xaxis | Hipp_L_Ant | -2.24 | 0.046119 | -0.38 |
| Hipp_L_Post2 | R_CB8 | -2.09 | 0.046119 | -0.36 |
| Hipp_L_Post2 | R_CB38 | -1.99 | 0.048173 | -0.34 |
| Hipp_L_Post1 | R_Dorsal_boundary_Zaxis | 2.25 | 0.048173 | 0.39 |

**Supplementary Table 4**. Cerebellar-hippocampal regions showing significant correlations in cortical FC with increased 17β-estradiol levels.

| **FC Seed Regions** | | **T(126)** | **pFDR** | **Cohen's d** |
|---|---|---|---|---|
| R_CB36 | Hipp_L_Ant_2 | 3.51 | 0.0147 | 0.63 |
| Hipp_L_Ant_2 | R_CB36 | 3.51 | 0.0147 | 0.63 |
| R_CB35 | Hipp_L_Ant_3_Rev2 | 3.25 | 0.0147 | 0.58 |
| Hipp_L_Ant_2 | R_Dorsal_boundary_Zaxis | 3.15 | 0.0147 | 0.56 |
| Hipp_L_Ant_2 | R_CB30 | 3.07 | 0.0147 | 0.55 |
| R_CB36 | Hipp_L_Ant_1 | 3.12 | 0.0147 | 0.56 |
| Hipp_L_Ant_2 | R_CB8 | 2.74 | 0.0147 | 0.49 |
| Hipp_L_Ant_3_Rev2 | R_CB35 | 3.25 | 0.0147 | 0.58 |
| R_CB35 | Hipp_L_Ant_1 | 2.88 | 0.0147 | 0.51 |
| R_Dorsal_boundary_Zaxis | Hipp_L_Ant_2 | 3.15 | 0.0147 | 0.56 |
| Hipp_L_Ant_1 | R_CB36 | 3.12 | 0.0217 | 0.56 |
| Hipp_L_Ant_1 | R_CB35 | 2.88 | 0.0217 | 0.51 |
| R_CB30 | Hipp_L_Ant_2 | 3.07 | 0.0231 | 0.55 |
| R_CB38 | Hipp_L_Ant | 2.81 | 0.0231 | 0.50 |
| Hipp_L_Ant_1 | R_Medial_boundary_Xaxis | 2.65 | 0.0243 | 0.47 |
| R_CB38 | Hipp_L_Post3 | 2.62 | 0.0243 | 0.47 |
| R_CB38 | Hipp_L_Ant_1 | 2.58 | 0.0253 | 0.46 |
| Hipp_L_Ant_1 | R_CB38 | 2.58 | 0.0253 | 0.46 |
| Hipp_L_Ant_2 | R_CB31 | 2.38 | 0.0253 | 0.42 |
| R_CB8 | Hipp_L_Ant_2 | 2.74 | 0.0253 | 0.49 |
| Hipp_L_Ant | R_CB38 | 2.81 | 0.0253 | 0.50 |
| Hipp_L_Ant | R_CB36 | 2.34 | 0.0253 | 0.42 |
| Hipp_L_Ant | R_CB18 | 2.23 | 0.0258 | 0.40 |
| Hipp_L_Ant | R_CB7 | 2.22 | 0.0258 | 0.40 |
| Hipp_L_Ant | R_Dorsal_boundary_Zaxis | 2.21 | 0.0285 | 0.39 |
| Hipp_L_Ant | R_Medial_boundary_Xaxis | 2.14 | 0.0285 | 0.38 |
| R_CB36 | Hipp_L_Post3 | 2.35 | 0.0363 | 0.42 |
| R_CB36 | Hipp_L_Ant | 2.34 | 0.0363 | 0.42 |
| R_CB37 | Hipp_L_Post1 | -2.65 | 0.0363 | -0.47 |
| Hipp_L_Ant_2 | R_CB15 | 2.03 | 0.0363 | 0.36 |
| R_CB35 | Hipp_L_Post3 | 2.18 | 0.0363 | 0.39 |
| Hipp_L_Ant_1 | R_Dorsal_boundary_Zaxis | 2.07 | 0.0363 | 0.37 |
| Hipp_L_Ant_1 | R_CB14 | 2.06 | 0.0431 | 0.37 |
| Hipp_L_Ant_1 | R_CB31 | 2.06 | 0.0431 | 0.37 |
| Hipp_L_Ant_1 | R_CB30 | 2.06 | 0.0431 | 0.37 |
| R_Dorsal_boundary_Zaxis | Hipp_L_Ant | 2.21 | 0.0431 | 0.39 |
| R_Dorsal_boundary_Zaxis | Hipp_L_Ant_1 | 2.07 | 0.0431 | 0.37 |
| R_CB36 | Hipp_L_Ant_3_Rev2 | 1.99 | 0.0431 | 0.35 |
| Hipp_L_Post3 | R_CB38 | 2.62 | 0.0439 | 0.47 |
| Hipp_L_Post3 | R_CB18 | 2.51 | 0.0439 | 0.45 |

| | | | | |
|---|---|---|---|---|
| Hipp_L_Post3 | R_CB36 | 2.35 | 0.0462 | 0.42 |
| R_CB30 | Hipp_L_Ant_1 | 2.06 | 0.0462 | 0.37 |
| R_CB18 | Hipp_L_Post3 | 2.51 | 0.0469 | 0.45 |
| R_CB18 | Hipp_L_Ant | 2.23 | 0.0469 | 0.40 |
| Hipp_L_Post3 | R_CB35 | 2.18 | 0.0469 | 0.39 |
| R_CB31 | Hipp_L_Ant_2 | 2.38 | 0.0469 | 0.42 |
| R_CB31 | Hipp_L_Ant_1 | 2.06 | 0.0469 | 0.37 |
| R_Medial_boundary_Xaxis | Hipp_L_Ant_1 | 2.65 | 0.0469 | 0.47 |
| Hipp_L_Ant_3_Rev2 | R_CB36 | 1.99 | 0.0469 | 0.35 |
| Hipp_L_Ant_3_Rev2 | R_Medial_boundary_Xaxis | 1.98 | 0.0469 | 0.35 |
| Hipp_L_Post1 | R_CB37 | -2.65 | 0.0474 | -0.47 |
| R_CB7 | Hipp_L_Ant | 2.22 | 0.0474 | 0.40 |
| R_CB15 | Hipp_L_Ant_2 | 2.03 | 0.0494 | 0.36 |
| R_Medial_boundary_Xaxis | Hipp_L_Ant | 2.14 | 0.0494 | 0.38 |
| R_Medial_boundary_Xaxis | Hipp_L_Ant_3_Rev2 | 1.98 | 0.0494 | 0.35 |
| R_CB14 | Hipp_L_Ant_1 | 2.06 | 0.0494 | 0.37 |

**Supplementary Table 5**. Cerebellar-hippocampal regions showing significant correlations in cortical FC with increased progesterone levels.

| FC Seed Regions | | T(128) | pFDR | Cohen's d |
|---|---|---|---|---|
| R_CB37 | Hipp_L_Post1 | -6.43 | 0.0000 | -1.14 |
| R_CB36 | Hipp_L_Post1 | -6.14 | 0.0000 | -1.09 |
| Hipp_L_Post1 | R_CB37 | -6.43 | 0.0000 | -1.14 |
| R_Dorsal_boundary_Zaxis | Hipp_L_Ant_1 | -5.81 | 0.0000 | -1.03 |
| Hipp_L_Post1 | R_CB36 | -6.14 | 0.0000 | -1.09 |
| R_CB30 | Hipp_L_Post2 | -5.92 | 0.0000 | -1.05 |
| R_CB38 | Hipp_L_Post1 | -5.82 | 0.0000 | -1.03 |
| Hipp_L_Post2 | R_CB30 | -5.92 | 0.0000 | -1.05 |
| Hipp_L_Ant_1 | R_Dorsal_boundary_ | -5.81 | 0.0000 | -1.03 |
| Hipp_L_Post1 | R_CB38 | -5.82 | 0.0000 | -1.03 |
| R_CB5 | Hipp_L_Post1 | -5.36 | 0.0000 | -0.95 |
| R_CB39 | Hipp_L_Ant_1 | -5.39 | 0.0000 | -0.95 |
| Hipp_L_Ant_1 | R_CB39 | -5.39 | 0.0000 | -0.95 |
| Hipp_L_Post1 | R_CB5 | -5.36 | 0.0000 | -0.95 |
| R_CB9 | Hipp_L_Ant_1 | -4.88 | 0.0000 | -0.86 |
| R_CB4 | Hipp_L_Post1 | -4.71 | 0.0000 | -0.83 |
| Hipp_L_Ant_1 | R_CB9 | -4.88 | 0.0000 | -0.86 |
| R_CB7 | Hipp_L_Post2 | -4.66 | 0.0000 | -0.82 |
| Hipp_L_Post1 | R_CB4 | -4.71 | 0.0000 | -0.83 |
| R_CB9 | Hipp_L_Post3 | -4.40 | 0.0000 | -0.78 |
| Hipp_L_Post2 | R_CB7 | -4.66 | 0.0000 | -0.82 |
| R_Dorsal_boundary_Zaxis | Hipp_L_Post3 | -4.22 | 0.0000 | -0.75 |
| Hipp_L_Post3 | R_CB9 | -4.40 | 0.0000 | -0.78 |

| | | | | |
|---|---|---|---|---|
| R_CB31 | Hipp_L_Post2 | -4.04 | 0.0001 | -0.71 |
| R_Rostral_boundary_Yaxis | Hipp_L_Post2 | -4.28 | 0.0000 | -0.76 |
| Hipp_L_Post2 | R_Rostral_boundary | -4.28 | 0.0000 | -0.76 |
| R_Dorsal_boundary_Zaxis | Hipp_L_Ant_2 | -4.01 | 0.0001 | -0.71 |
| Hipp_L_Post3 | R_Dorsal_boundary_ | -4.22 | 0.0000 | -0.75 |
| Hipp_L_Post2 | R_CB31 | -4.04 | 0.0001 | -0.71 |
| R_Ventral_boundary_Zaxis | Hipp_L_Ant_1 | 3.98 | 0.0001 | 0.70 |
| R_CB7 | Hipp_L_Post1 | -3.9 | 0.0002 | -0.69 |
| R_Dorsal_boundary_Zaxis | Hipp_L_Ant_3_Rev2 | -3.77 | 0.0002 | -0.67 |
| R_Dorsal_boundary_Zaxis | Hipp_L_Post2 | -3.73 | 0.0003 | -0.66 |
| Hipp_L_Ant_2 | R_Dorsal_boundary_ | -4.01 | 0.0001 | -0.71 |
| R_CB9 | Hipp_L_Post1 | -3.61 | 0.0004 | -0.64 |
| Hipp_L_Ant_1 | R_Ventral_boundary | 3.98 | 0.0001 | 0.70 |
| Hipp_L_Post1 | R_CB7 | -3.90 | 0.0002 | -0.69 |
| R_CB19 | Hipp_L_Post3 | 3.67 | 0.0003 | 0.65 |
| R_Medial_boundary_Xaxis | Hipp_L_Ant_2 | -3.59 | 0.0005 | -0.63 |
| R_CB39 | Hipp_L_Ant_2 | -3.64 | 0.0004 | -0.64 |
| Hipp_L_Post2 | R_Dorsal_boundary_ | -3.73 | 0.0003 | -0.66 |
| R_CB9 | Hipp_L_Ant_3_Rev2 | -3.46 | 0.0007 | -0.61 |
| R_CB9 | Hipp_L_Ant_2 | -3.46 | 0.0007 | -0.61 |
| R_CB26 | Hipp_L_Post2 | -3.51 | 0.0006 | -0.62 |
| R_CB39 | Hipp_L_Ant_3_Rev2 | -3.49 | 0.0007 | -0.62 |
| R_CB18 | Hipp_L_Post1 | -3.50 | 0.0006 | -0.62 |
| Hipp_L_Post3 | R_CB19 | 3.67 | 0.0003 | 0.65 |
| Hipp_L_Post1 | R_CB9 | -3.61 | 0.0004 | -0.64 |
| Hipp_L_Post2 | R_CB26 | -3.51 | 0.0006 | -0.62 |
| Hipp_L_Post1 | R_CB18 | -3.5 | 0.0006 | -0.62 |
| Hipp_L_Ant_2 | R_CB39 | -3.64 | 0.0004 | -0.64 |
| R_Ventral_boundary_Zaxis | Hipp_L_Ant_3_Rev2 | 3.46 | 0.0007 | 0.61 |
| R_Caudal_boundary_Yaxis | Hipp_L_Post2 | -3.29 | 0.0013 | -0.58 |
| Hipp_L_Ant_2 | R_Medial_boundary_ | -3.59 | 0.0005 | -0.63 |
| Hipp_L_Ant_3_Rev2 | R_Dorsal_boundary_ | -3.77 | 0.0002 | -0.67 |
| R_CB35 | Hipp_L_Ant_3_Rev2 | -3.21 | 0.0017 | -0.57 |
| R_CB19 | Hipp_L_Post2 | 3.23 | 0.0016 | 0.57 |
| Hipp_L_Ant_2 | R_CB9 | -3.46 | 0.0007 | -0.61 |
| R_Ventral_boundary_Zaxis | Hipp_L_Post3 | 3.27 | 0.0014 | 0.58 |
| Hipp_L_Post2 | R_Caudal_boundary_ | -3.29 | 0.0013 | -0.58 |
| R_CB4 | Hipp_L_Post2 | -3.14 | 0.0021 | -0.56 |
| Hipp_L_Post2 | R_CB19 | 3.23 | 0.0016 | 0.57 |
| Hipp_L_Post3 | R_Ventral_boundary | 3.27 | 0.0014 | 0.58 |
| Hipp_L_Ant_3_Rev2 | R_CB39 | -3.49 | 0.0007 | -0.62 |
| Hipp_L_Ant_3_Rev2 | R_CB9 | -3.46 | 0.0007 | -0.61 |
| Hipp_L_Ant_3_Rev2 | R_Ventral_boundary | 3.46 | 0.0007 | 0.61 |
| Hipp_L_Post2 | R_CB4 | -3.14 | 0.0021 | -0.56 |
| R_Medial_boundary_Xaxis | Hipp_L_Post2 | -2.97 | 0.0036 | -0.53 |

| | | | | |
|---|---|---|---|---|
| R_Caudal_boundary_Yaxis | Hipp_L_Ant_1 | -2.86 | 0.0050 | -0.51 |
| Hipp_L_Post2 | R_Medial_boundary_ | -2.97 | 0.0036 | -0.53 |
| Hipp_L_Ant_3_Rev2 | R_CB35 | -3.21 | 0.0017 | -0.57 |
| R_Dorsal_boundary_Zaxis | Hipp_L_Ant | -2.78 | 0.0063 | -0.49 |
| R_Medial_boundary_Xaxis | Hipp_L_Ant | -2.6 | 0.0105 | -0.46 |
| R_Caudal_boundary_Yaxis | Hipp_L_Post3 | -2.59 | 0.0107 | -0.46 |
| R_CB37 | Hipp_L_Ant_3_Rev2 | -2.56 | 0.0116 | -0.45 |
| R_CB39 | Hipp_L_Post3 | -2.6 | 0.0103 | -0.46 |
| R_CB35 | Hipp_L_Post1 | -2.56 | 0.0116 | -0.45 |
| Hipp_L_Ant_1 | R_Caudal_boundary_ | -2.86 | 0.0050 | -0.51 |
| R_CB14 | Hipp_L_Ant_1 | -2.61 | 0.0102 | -0.46 |
| R_CB8 | Hipp_L_Post1 | -2.41 | 0.0175 | -0.43 |
| Hipp_L_Post1 | R_CB35 | -2.56 | 0.0116 | -0.45 |
| Hipp_L_Post3 | R_CB39 | -2.60 | 0.0103 | -0.46 |
| Hipp_L_Post3 | R_Caudal_boundary_ | -2.59 | 0.0107 | -0.46 |
| R_Medial_boundary_Xaxis | Hipp_L_Ant_3_Rev2 | -2.32 | 0.0218 | -0.41 |
| Hipp_L_Ant_1 | R_CB14 | -2.61 | 0.0102 | -0.46 |
| R_CB30 | Hipp_L_Post1 | -2.39 | 0.0183 | -0.42 |
| R_CB18 | Hipp_L_Ant_1 | 2.31 | 0.0225 | 0.41 |
| R_CB15 | Hipp_L_Post3 | 2.30 | 0.0233 | 0.41 |
| R_Caudal_boundary_Yaxis | Hipp_L_Post1 | -2.22 | 0.0280 | -0.39 |
| R_CB35 | Hipp_L_Ant | -2.22 | 0.0281 | -0.39 |
| R_CB5 | Hipp_L_Post2 | -2.25 | 0.0263 | -0.40 |
| Hipp_L_Post1 | R_CB8 | -2.41 | 0.0175 | -0.43 |
| Hipp_L_Post1 | R_CB30 | -2.39 | 0.0183 | -0.42 |
| R_CB39 | Hipp_L_Ant | -2.19 | 0.0302 | -0.39 |
| R_Caudal_boundary_Yaxis | Hipp_L_Ant | -2.14 | 0.0339 | -0.38 |
| R_CB36 | Hipp_L_Post2 | -2.16 | 0.0328 | -0.38 |
| R_CB19 | Hipp_L_Post1 | -2.23 | 0.0274 | -0.39 |
| R_Rostral_boundary_Yaxis | Hipp_L_Post1 | -2.28 | 0.0243 | -0.40 |
| Hipp_L_Post1 | R_Rostral_boundary | -2.28 | 0.0243 | -0.40 |
| Hipp_L_Ant | R_Dorsal_boundary_ | -2.78 | 0.0063 | -0.49 |
| Hipp_L_Post1 | R_CB19 | -2.23 | 0.0274 | -0.39 |
| Hipp_L_Post1 | R_Caudal_boundary_ | -2.22 | 0.0280 | -0.39 |
| R_CB9 | Hipp_L_Ant | -2.05 | 0.0425 | -0.36 |
| Hipp_L_Post2 | R_CB5 | -2.25 | 0.0263 | -0.40 |
| Hipp_L_Ant_3_Rev2 | R_CB37 | -2.56 | 0.0116 | -0.45 |
| Hipp_L_Post3 | R_CB15 | 2.30 | 0.0233 | 0.41 |
| R_Medial_boundary_Xaxis | Hipp_L_Ant_1 | -2.02 | 0.0456 | -0.36 |
| R_CB26 | Hipp_L_Post1 | -2.00 | 0.0478 | -0.35 |
| R_CB8 | Hipp_L_Post3 | -2.01 | 0.0461 | -0.36 |
| Hipp_L_Post2 | R_CB36 | -2.16 | 0.0328 | -0.38 |
| Hipp_L_Ant | R_Medial_boundary_ | -2.60 | 0.0105 | -0.46 |
| Hipp_L_Ant_1 | R_CB18 | 2.31 | 0.0225 | 0.41 |
| R_Ventral_boundary_Zaxis | Hipp_L_Ant_2 | 2.05 | 0.0428 | 0.36 |

| | | | | |
|---|---|---|---|---|
| R_CB7 | Hipp_L_Ant_1 | -2.05 | 0.0424 | -0.36 |
| R_CB19 | Hipp_L_Ant_2 | -1.99 | 0.0492 | -0.35 |
| Hipp_L_Post1 | R_CB26 | -2.00 | 0.0478 | -0.35 |
| Hipp_L_Ant_3_Rev2 | R_Medial_boundary_ | -2.32 | 0.0218 | -0.41 |
| Hipp_L_Post3 | R_CB8 | -2.01 | 0.0461 | -0.36 |
| Hipp_L_Ant_1 | R_CB7 | -2.05 | 0.0424 | -0.36 |
| Hipp_L_Ant_1 | R_Medial_boundary | -2.02 | 0.0456 | -0.36 |
| Hipp_L_Ant_2 | R_Ventral_boundary | 2.05 | 0.0428 | 0.36 |
| Hipp_L_Ant_2 | R_CB19 | -1.99 | 0.0492 | -0.35 |
| Hipp_L_Ant | R_CB35 | -2.22 | 0.0281 | -0.39 |
| Hipp_L_Ant | R_CB39 | -2.19 | 0.0302 | -0.39 |
| Hipp_L_Ant | R_Caudal_boundary_ | -2.14 | 0.0339 | -0.38 |
| Hipp_L_Ant | R_CB9 | -2.05 | 0.0425 | -0.36 |
| R_CB32 | Hipp_L_Post2 | -5.77 | 0.0000 | -1.02 |
| R_CB29 | Hipp_L_Post3 | -5.81 | 0.0000 | -1.03 |
| Hipp_L_Post3 | R_CB29 | -5.81 | 0.0000 | -1.03 |
| Hipp_L_Post2 | R_CB32 | -5.77 | 0.0000 | -1.02 |
| R_CB33 | Hipp_L_Post3 | -5.28 | 0.0000 | -0.93 |
| R_CB33 | Hipp_L_Ant_1 | -5.22 | 0.0000 | -0.92 |
| R_CB29 | Hipp_L_Post2 | -5.15 | 0.0000 | -0.91 |
| Hipp_L_Post3 | R_CB33 | -5.28 | 0.0000 | -0.93 |
| R_CB2 | Hipp_L_Post3 | -4.99 | 0.0000 | -0.88 |
| Hipp_L_Ant_1 | R_CB33 | -5.22 | 0.0000 | -0.92 |
| Hipp_L_Post2 | R_CB29 | -5.15 | 0.0000 | -0.91 |
| R_CB6 | Hipp_L_Post1 | -4.73 | 0.0000 | -0.84 |
| R_CB33 | Hipp_L_Post2 | -4.76 | 0.0000 | -0.84 |
| R_CB27 | Hipp_L_Post2 | -4.66 | 0.0000 | -0.82 |
| Hipp_L_Post3 | R_CB2 | -4.99 | 0.0000 | -0.88 |
| R_CB27 | Hipp_L_Ant_2 | -4.59 | 0.0000 | -0.81 |
| R_CB2 | Hipp_L_Ant_1 | -4.64 | 0.0000 | -0.82 |
| R_CB32 | Hipp_L_Ant_2 | -4.54 | 0.0000 | -0.80 |
| Hipp_L_Post2 | R_CB33 | -4.76 | 0.0000 | -0.84 |
| Hipp_L_Post1 | R_CB6 | -4.73 | 0.0000 | -0.84 |
| R_CB2 | Hipp_L_Post1 | -4.46 | 0.0000 | -0.79 |
| Hipp_L_Post2 | R_CB27 | -4.66 | 0.0000 | -0.82 |
| Hipp_L_Ant_1 | R_CB2 | -4.64 | 0.0000 | -0.82 |
| R_CB22 | Hipp_L_Post2 | -4.28 | 0.0000 | -0.76 |
| R_CB27 | Hipp_L_Post1 | -4.17 | 0.0001 | -0.74 |
| Hipp_L_Post1 | R_CB2 | -4.46 | 0.0000 | -0.79 |
| Hipp_L_Ant_2 | R_CB27 | -4.59 | 0.0000 | -0.81 |
| Hipp_L_Ant_2 | R_CB32 | -4.54 | 0.0000 | -0.80 |
| R_CB22 | Hipp_L_Ant_2 | -4.14 | 0.0001 | -0.73 |
| Hipp_L_Post2 | R_CB22 | -4.28 | 0.0000 | -0.76 |
| R_CB32 | Hipp_L_Ant_1 | -3.97 | 0.0001 | -0.70 |
| R_CB23 | Hipp_L_Ant_2 | -4.04 | 0.0001 | -0.71 |

| | | | | |
|---|---|---|---|---|
| R_CB22 | Hipp_L_Post1 | -3.97 | 0.0001 | -0.70 |
| Hipp_L_Post1 | R_CB27 | -4.17 | 0.0001 | -0.74 |
| R_CB27 | Hipp_L_Post3 | -3.84 | 0.0002 | -0.68 |
| R_CB28 | Hipp_L_Post2 | -3.78 | 0.0002 | -0.67 |
| R_CB23 | Hipp_L_Post1 | -3.82 | 0.0002 | -0.68 |
| Hipp_L_Ant_2 | R_CB22 | -4.14 | 0.0001 | -0.73 |
| Hipp_L_Post1 | R_CB22 | -3.97 | 0.0001 | -0.70 |
| R_CB29 | Hipp_L_Post1 | -3.69 | 0.0003 | -0.65 |
| Hipp_L_Ant_2 | R_CB23 | -4.04 | 0.0001 | -0.71 |
| Hipp_L_Ant_1 | R_CB32 | -3.97 | 0.0001 | -0.70 |
| R_CB29 | Hipp_L_Ant_1 | -3.57 | 0.0005 | -0.63 |
| Hipp_L_Post1 | R_CB23 | -3.82 | 0.0002 | -0.68 |
| Hipp_L_Post2 | R_CB28 | -3.78 | 0.0002 | -0.67 |
| R_CB16 | Hipp_L_Post1 | -3.55 | 0.0005 | -0.63 |
| Hipp_L_Post3 | R_CB27 | -3.84 | 0.0002 | -0.68 |
| R_CB21 | Hipp_L_Post1 | -3.55 | 0.0005 | -0.63 |
| Hipp_L_Post1 | R_CB29 | -3.69 | 0.0003 | -0.65 |
| R_CB28 | Hipp_L_Post1 | -3.38 | 0.0010 | -0.60 |
| R_CB13 | Hipp_L_Post1 | -3.42 | 0.0008 | -0.60 |
| R_CB28 | Hipp_L_Ant_1 | -3.34 | 0.0011 | -0.59 |
| Hipp_L_Post1 | R_CB21 | -3.55 | 0.0005 | -0.63 |
| Hipp_L_Post1 | R_CB16 | -3.55 | 0.0005 | -0.63 |
| R_CB24 | Hipp_L_Post3 | -3.31 | 0.0012 | -0.59 |
| Hipp_L_Post1 | R_CB13 | -3.42 | 0.0008 | -0.60 |
| Hipp_L_Ant_1 | R_CB29 | -3.57 | 0.0005 | -0.63 |
| Hipp_L_Post1 | R_CB28 | -3.38 | 0.0010 | -0.60 |
| R_CB17 | Hipp_L_Post3 | 3.27 | 0.0014 | 0.58 |
| R_CB22 | Hipp_L_Ant_1 | -3.15 | 0.0020 | -0.56 |
| R_CB23 | Hipp_L_Post3 | -3.19 | 0.0018 | -0.56 |
| R_CB28 | Hipp_L_Ant_2 | -3.12 | 0.0022 | -0.55 |
| R_CB24 | Hipp_L_Post1 | -3.13 | 0.0022 | -0.55 |
| R_CB23 | Hipp_L_Post2 | -3.1 | 0.0024 | -0.55 |
| R_CB22 | Hipp_L_Post3 | -3.01 | 0.0032 | -0.53 |
| Hipp_L_Post3 | R_CB24 | -3.31 | 0.0012 | -0.59 |
| Hipp_L_Post3 | R_CB17 | 3.27 | 0.0014 | 0.58 |
| Hipp_L_Ant_1 | R_CB28 | -3.34 | 0.0011 | -0.59 |
| Hipp_L_Post1 | R_CB24 | -3.13 | 0.0022 | -0.55 |
| R_CB2 | Hipp_L_Ant | -2.98 | 0.0035 | -0.53 |
| Hipp_L_Post3 | R_CB23 | -3.19 | 0.0018 | -0.56 |
| Hipp_L_Post2 | R_CB23 | -3.1 | 0.0024 | -0.55 |
| Hipp_L_Ant_1 | R_CB22 | -3.15 | 0.0020 | -0.56 |
| Hipp_L_Ant_2 | R_CB28 | -3.12 | 0.0022 | -0.55 |
| R_CB33 | Hipp_L_Ant_3_Rev2 | -2.75 | 0.0068 | -0.49 |
| Hipp_L_Post3 | R_CB22 | -3.01 | 0.0032 | -0.53 |
| R_CB27 | Hipp_L_Ant_1 | -2.67 | 0.0085 | -0.47 |

| | | | | |
|---|---|---|---|---|
| R_CB28 | Hipp_L_Ant | -2.67 | 0.0086 | -0.47 |
| R_CB2 | Hipp_L_Post2 | -2.72 | 0.0075 | -0.48 |
| R_CB6 | Hipp_L_Ant_2 | -2.68 | 0.0083 | -0.47 |
| R_CB33 | Hipp_L_Ant_2 | -2.58 | 0.0110 | -0.46 |
| R_CB2 | Hipp_L_Ant_2 | -2.58 | 0.0111 | -0.46 |
| Hipp_L_Post2 | R_CB2 | -2.72 | 0.0075 | -0.48 |
| R_CB24 | Hipp_L_Post2 | -2.5 | 0.0138 | -0.44 |
| R_CB32 | Hipp_L_Post1 | -2.41 | 0.0174 | -0.43 |
| R_CB28 | Hipp_L_Ant_3_Rev2 | -2.36 | 0.0199 | -0.42 |
| Hipp_L_Ant_1 | R_CB27 | -2.67 | 0.0085 | -0.47 |
| Hipp_L_Post2 | R_CB24 | -2.5 | 0.0138 | -0.44 |
| Hipp_L_Ant_2 | R_CB6 | -2.68 | 0.0083 | -0.47 |
| R_CB32 | Hipp_L_Ant_3_Rev2 | -2.26 | 0.0257 | -0.40 |
| Hipp_L_Ant | R_CB2 | -2.98 | 0.0035 | -0.53 |
| R_CB28 | Hipp_L_Post3 | -2.21 | 0.0287 | -0.39 |
| Hipp_L_Ant_2 | R_CB33 | -2.58 | 0.0110 | -0.46 |
| Hipp_L_Ant_2 | R_CB2 | -2.58 | 0.0111 | -0.46 |
| Hipp_L_Ant_3_Rev2 | R_CB33 | -2.75 | 0.0068 | -0.49 |
| R_Lateral_boundary_Xaxis | Hipp_L_Ant_2 | -2.3 | 0.0232 | -0.41 |
| Hipp_L_Post1 | R_CB32 | -2.41 | 0.0174 | -0.43 |
| R_CB32 | Hipp_L_Post3 | -2.1 | 0.0378 | -0.37 |
| R_CB17 | Hipp_L_Post1 | -2.18 | 0.0313 | -0.39 |
| R_CB17 | Hipp_L_Ant_3_Rev2 | 2.13 | 0.0351 | 0.38 |
| R_Lateral_boundary_Xaxis | Hipp_L_Ant_1 | -2.07 | 0.0401 | -0.37 |
| Hipp_L_Post1 | R_CB17 | -2.18 | 0.0313 | -0.39 |
| R_CB21 | Hipp_L_Post3 | 2.06 | 0.0414 | 0.36 |
| Hipp_L_Ant | R_CB28 | -2.67 | 0.0086 | -0.47 |
| R_CB25 | Hipp_L_Post3 | -2.1 | 0.0377 | -0.37 |
| Hipp_L_Post3 | R_CB28 | -2.21 | 0.0287 | -0.39 |
| R_CB25 | Hipp_L_Post2 | 2.04 | 0.0431 | 0.36 |
| Hipp_L_Ant_2 | R_Lateral_boundary | -2.3 | 0.0232 | -0.41 |
| Hipp_L_Post2 | R_CB25 | 2.04 | 0.0431 | 0.36 |
| Hipp_L_Post3 | R_CB25 | -2.1 | 0.0377 | -0.37 |
| Hipp_L_Post3 | R_CB32 | -2.1 | 0.0378 | -0.37 |
| Hipp_L_Ant_3_Rev2 | R_CB28 | -2.36 | 0.0199 | -0.42 |
| Hipp_L_Post3 | R_CB21 | 2.06 | 0.0414 | 0.36 |
| Hipp_L_Ant_3_Rev2 | R_CB32 | -2.26 | 0.0257 | -0.40 |
| Hipp_L_Ant_1 | R_Lateral_boundary | -2.07 | 0.0401 | -0.37 |
| Hipp_L_Ant_3_Rev2 | R_CB17 | 2.13 | 0.0351 | 0.38 |
| R_CB11 | Hipp_L_Post1 | -5.2 | 0.0000 | -0.92 |
| R_CB1 | Hipp_L_Post1 | -5.2 | 0.0000 | -0.92 |
| Hipp_L_Post1 | R_CB11 | -5.2 | 0.0000 | -0.92 |
| Hipp_L_Post1 | R_CB1 | -5.2 | 0.0000 | -0.92 |
| R_CB1 | Hipp_L_Post2 | -4.09 | 0.0001 | -0.72 |
| R_CB11 | Hipp_L_Post2 | -4.09 | 0.0001 | -0.72 |

| | | | | |
|---|---|---|---|---|
| Hipp_L_Post2 | R_CB11 | -4.09 | 0.0001 | -0.72 |
| Hipp_L_Post2 | R_CB1 | -4.09 | 0.0001 | -0.72 |
| R_CB11 | Hipp_L_Post3 | -3.67 | 0.0004 | -0.65 |
| R_CB1 | Hipp_L_Post3 | -3.67 | 0.0004 | -0.65 |
| Hipp_L_Post3 | R_CB11 | -3.67 | 0.0004 | -0.65 |
| Hipp_L_Post3 | R_CB1 | -3.67 | 0.0004 | -0.65 |
| R_CB11 | Hipp_L_Ant_1 | -2 | 0.0478 | -0.35 |
| R_CB1 | Hipp_L_Ant_1 | -2 | 0.0478 | -0.35 |
| Hipp_L_Ant_1 | R_CB11 | -2 | 0.0478 | -0.35 |
| Hipp_L_Ant_1 | R_CB1 | -2 | 0.0478 | -0.35 |

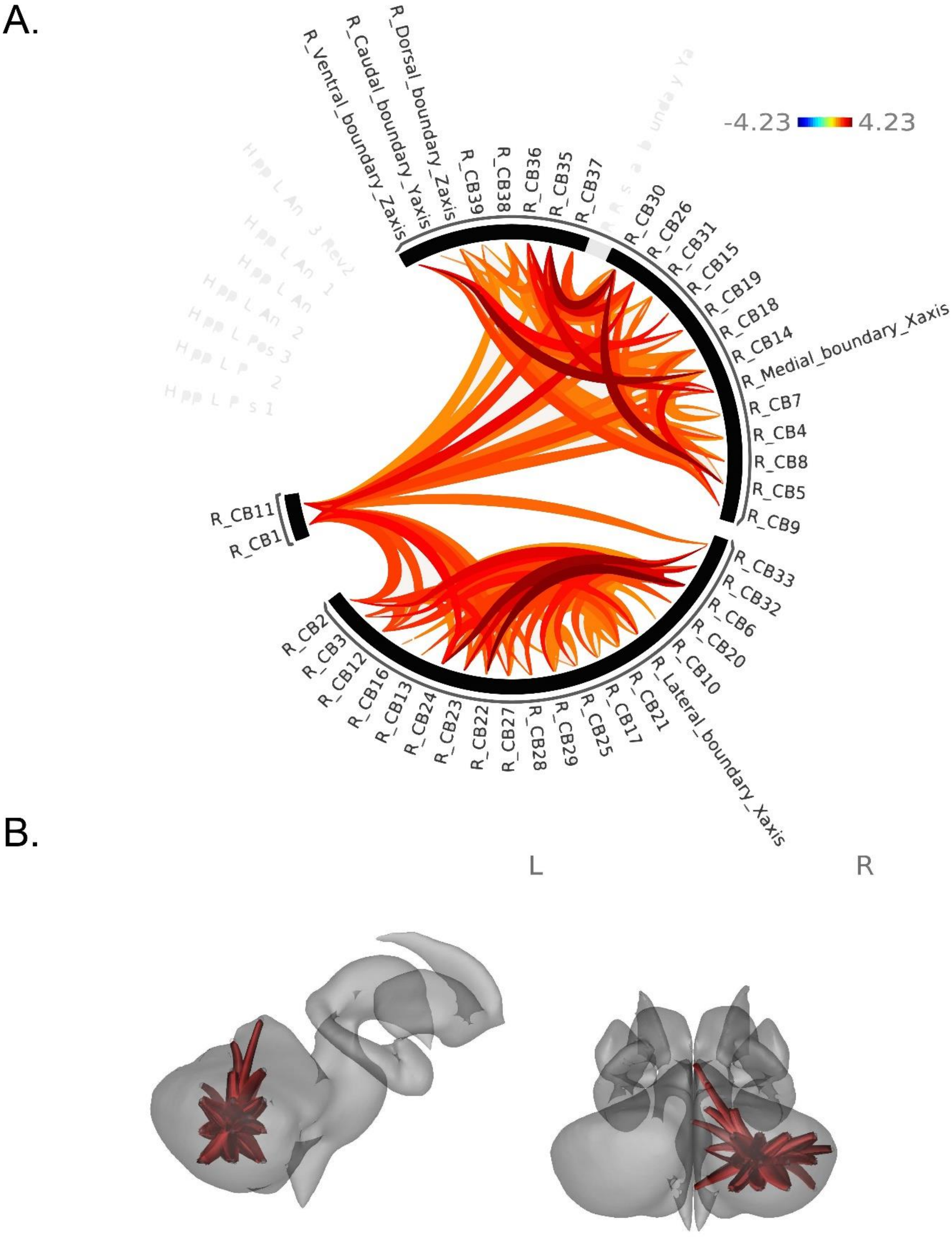


**Supplementary Figure 2.** Patterns of cortical functional connectivity (FC) between ROIs in males as compared to females, when controlling for age. **A**. ROIs are shown in an FC ring where orange-red displays greater FC in males as compared to females. **B**. ROIs are shown on a subcortical model where red displays positive intracerebellar FC relationships between ROIs in males as compared to females.

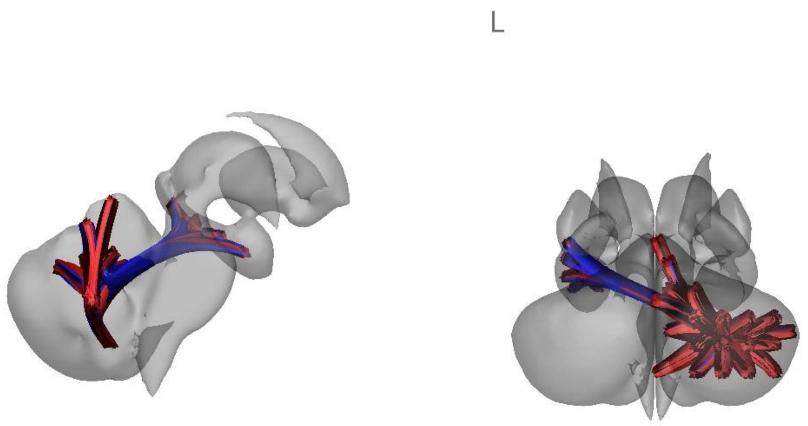


**Supplementary Figure 3**. Cerebellar and hippocampal regions showing significant within structure correlations in cortical FC with increased progesterone levels, when controlling for age. ROIs are shown on a subcortical model where blue represents negative FC relationships between ROIs with higher progesterone levels, while red displays positive FC relationships between ROIs with higher progesterone levels.

**Supplementary Table 6**. Select behavioral performances show negative relationships with age.

| Behavioral Measure | Main Effect of Age (pFDR) | Main Effect of Education Level (pFDR) | df |
|---|---|---|---|
| Symbol Span | 0.006* | 0.586 | 135 |
| Stroop Task | 0.242 | 0.642 | 126 |
| Pegboard Assembly | <0.001* | 0.691 | 135 |
| Shopping List Memory | 0.005* | 0.228 | 126 |
| Sequence Learning | 0.009* | 0.707 | 104 |
| Letter-Number Sequencing | <0.001* | 0.408 | 135 |

*Note.* * indicates significant p-value after FDR correction.*
